# The Signaling Pathway of a Bacterial Two-Component System Determined by cryo-EM

**DOI:** 10.64898/2026.10.12.758521

**Authors:** Prabin Karki, Carolina Hernandez, Joshua H. Mendez, William C. Budell, David Menendez, Shemiran Kanou, Sorayya Razavi, Pakhshan Espoukeh, Rotan Abuhashish, Dorina Bizhga, Shibom Basu, Tek Narsingh Malla, Peter Schwander, Emina A. Stojković, Marius Schmidt

**Affiliations:** Department of Physics and Astronomy, University of Wisconsin Milwaukee; Milwaukee, 53211, USA; New York Structural Biology Center; New York, 10027, USA; Department of Biology, Northeastern Illinois University; Chicago, 60625, USA; European Molecular Biology Laboratory (EMBL); Grenoble, 38043, France; Vaccine Research Center Electron Microscopy Unit, Frederick National Laboratory for Cancer Research; Frederick, 21701, USA; Department of Pharmaceutical and Administrative Sciences, School of Pharmacy, Concordia University of Wisconsin; Mequon, 53097, USA

## Abstract

Although two-component systems (TCSs) play ubiquitous roles in sensing signals for bacterial survival and virulence, a detailed structural map of the signal transduction pathway is lacking. Bacteriophytochromes (BphPs) are enzymatic photoreceptors with homologs in plants that regulate bacterial physiological responses via TCS signaling. Here, we employ cryo-EM to elucidate the structural basis of a light-regulated TCS by capturing the full-length BphP histidine kinase (HK) in complex with its downstream response regulator (RR). The RR binds as a dimer to the HK but contacts the enzyme only through one subunit. Light absorption perturbs the entire BphP structure, including the HK domain, which disrupts phosphorylation and phosphotransfer reactions and promote TCS complex dissociation. Thus, light controls signaling through both HK activity and reversible binding of the RR.

## Introduction

Two-component systems represent the most abundant signal transduction pathway in biology and are ubiquitous in bacteria (*1*). Composed of a sensor Histidine Kinase (HK) and its cognate response regulator (RR), they play essential roles in responding to environmental and intracellular cues (*2, 3*). In addition to their roles in cell metabolism, virulence and antibiotic resistance (*2*), TCSs are gaining attention in synthetic biology (*4, 5*) and antimicrobial drug development (*6*). Although many TCS pathways have been functionally characterized, limited structural knowledge exists to date regarding the interaction and formation of the HK-RR complex, which is essential to the generation and to the regulation of the signal. The crystal structure of the CheY RR has been solved more than 30 years ago (*7*). Crystal structures of HK-RR complexes were published in 2009 and 2016 (*8–10*). In these structures a cytosolic fragment of the HK was crystallized in complex with the RR. A symmetric, cross-like structure is displayed where the HK lobes alter with RR monomers. Since then, this structural topology was assumed to represent a canonical HK-RR complex. In typical TCSs, HKs act as donors of phosphate groups to their respective RRs, leading to specific output responses. The HK is frequently a membrane-associated protein that shares fundamental signaling components: a diverse sensing domain (light or ligand sensing), a signal-transducing domain, and a highly conserved kinase composed of a histidine phosphotransfer (DHp) domain, as well as an ATP/ADP-binding catalytic (CA) domain. In this study, we present cryo-EM structures of a classical, light-sensing HK bound to its cognate RR. The HK domain is part of a red-light sensing bacterial phytochrome (BphP) found in the cytoplasm of the soil-dwelling, aerobic mesophile, myxobacterium *Allostigmatella aurantiaca* (*11*). BphPs are part of TCS pathways across different bacterial kingdoms and members of a larger phytochrome family, originally discovered in plants. We show that the interaction of the full-length BphP with the RR is asymmetric, with only one subunit of the RR dimer interacting with the HK. Furthermore, the HK-RR complex is disrupted by absorption of a single photon by the light-sensing domain, that induces a 180° rotation of the HK domain (*12, 13*) and a downstream dissociation of the RR. This mechanism is corroborated by *in-vitro* phosphotransfer assays under different light conditions. The dynamic mapping of the TCS pathway by cryo-EM provides direct insight into the structural plasticity of HK interaction with the RR and how light and/or any other environmental signal are perceived by bacteria.

Myxobacteria are fruiting body forming eubacteria (*14*) where light stimulates the growth of their fruiting bodies (*15*) as well as the biosynthesis of secondary metabolites, many of which exhibit antibiotic or anticancer properties (*16*). Myxobacteria are also of considerable interest in agriculture because of their potential to suppress and control plant pathogens (*17–19*). *A. aurantiaca* features two enzymatic bacteriophytochromes (BphPs). Both BphPs include a C-terminal HK (Fig. 1 a,b) and are implicated in fruiting body formation via a canonical TCS (*11, 15*). BphPs are part of a large family of phytochromes, found in plants, diatoms and fungi (*20*). In plants, phytochromes regulate important responses such as shade avoidance and seed germination. BphPs are homodimers that consist of PAS (Per-Arnt-Sim), GAF (cGMP phosphodiesterase/adenylyl cyclase/FhIA), and PHY (phytochrome-specific GAF-related) domains that form the photosensory core module (PCM) which senses red and far-red light through a covalently bound biliverdin (BV) chromophore (Fig. 1 a-c). The isomerization within BV initiates the transition of the protein from the red-absorbing Pr to the far-red absorbing Pfr state (Fig. 1 c). Notably, BphP signal response occurs sequentially by activation of only one BV which will generate a signaling heterodimer (Fig. 1 a, c) where one monomer is in the Pr state the other in the Pfr state (*21*). This Pr/Pfr heterodimer is an intermediate in BphP photocycle (Fig. 1 c) which can be generated and observed on a millisecond time-scale by time-resolved cryo-EM (*13*). The transition from the dark-adapted Pr/Pr homodimer to light-triggered Pr/Pfr heterodimer involves a ∼180° rotation of the entire HK domain (*13*). In the process, the linker helices which form a coiled coil partially unwind. As a result, the HK lobes interchange position and cross the dimer interface (Fig. 1 a). Absorption of a second photon is associated with a further unwinding of the linker helices and a rotation of the HK, which results in an all-parallel linker helix configuration (*21*). These findings suggest that the unwinding of linker helices and the concomitant HK rotation provide an additional, and potentially dominant, mechanism for controlling signal transmission to the RR.

**Fig 1.**
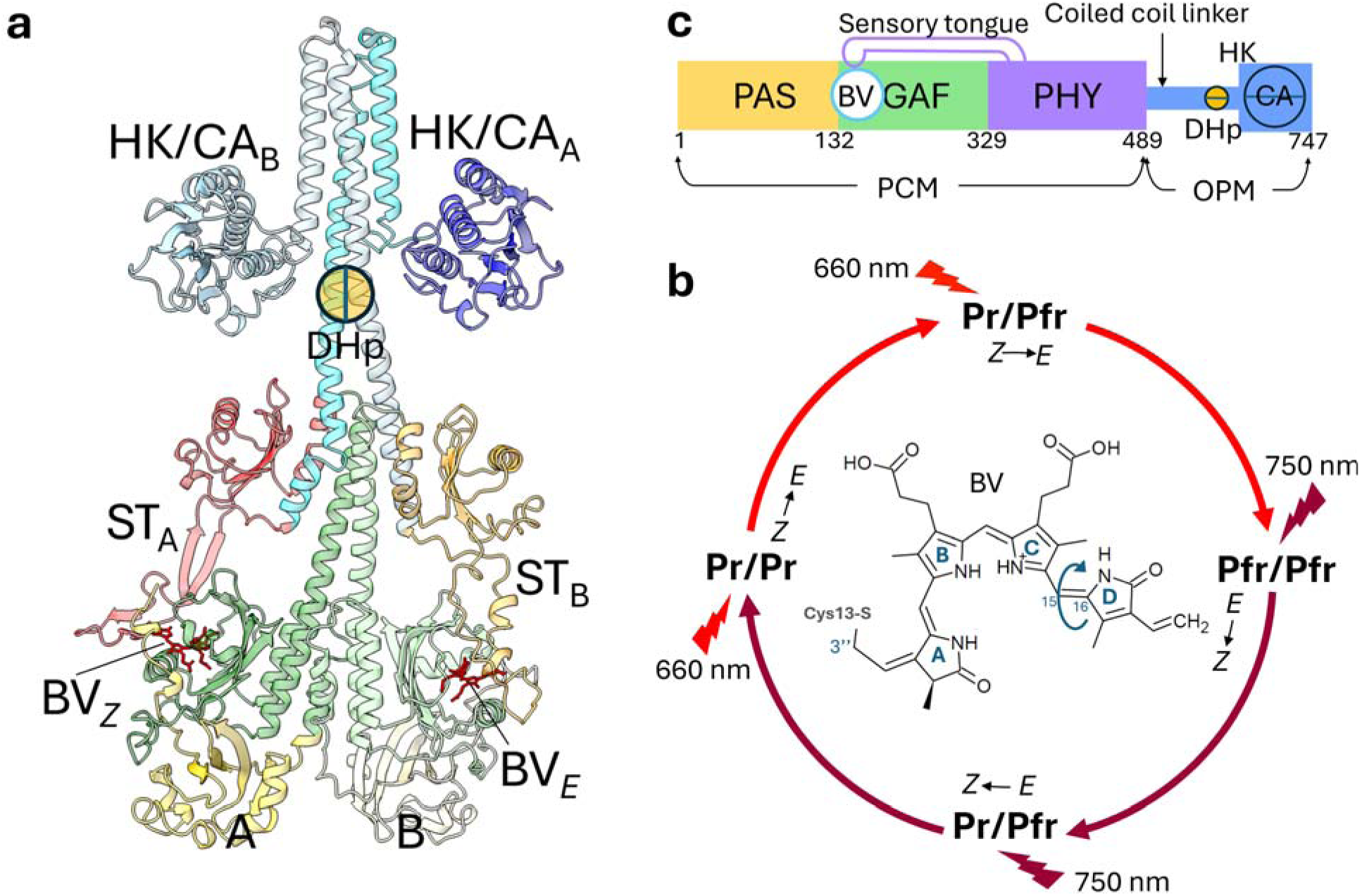
Structure and function of SaBphP2, a light sensing HK. **(a)** The structure of SaBphP2 from *A. aurantiaca* in the Pr/Pfr heterodimeric state (*21*). The biliverdins are marked BVz (ring-D in the Z-configuration) and BV*_E_*(ring-D in the E-configuration). The sensory tongues (ST) of protomers A and B are marked. ST_A_ forms a β-sheet indicative of the Pr state and ST_B_ an α-helix characteristic of the Pfr state. The HK catalytic domains (CA) are associated with the HK lobes, the phophotransfer domain (DHp) consists of a stretch of the linker around His523. **(b)** Domain organization of SaBphP2 mapped onto the amino acid sequence. The photosensory core module (PCM), the catalytic domain (CA) and the location of the DHp and HK domains are marked. **(c)** The SaBphP2 photocycle. The first half of the photocycle from Pr to Pfr is initiated by red light (660 nm, red flash), the second half from Pfr to Pr by far-red (750 nm, far-red flash) light. The Pr/Pfr heterodimer is a distinct photocycle intermediate. The chemical structure of the biliverdin (BV) chromophore is shown in the center. Absorption of a photon causes ring-D to isomerize from the *Z* to the *E* configuration by rotation about the 15-16 double bond.

## Results

### A. aurantiaca TCS - The SaBphP2– RR complex

Using cryo-EM, we map the light-regulated two component signaling (TCS) pathway in *A. aurantiaca*. To do so, we determined the structure of an intact HK as a part of a BphP. Here, we investigate the structure of SaBphP2 (*11*) which is the second BphP in *A. auratiaca*. The downstream response regulator associated with SaBphP2 is denoted as RR_SaBphP2_ and shares sequence similarity to CheY-like RRs (*2, 7*). In the presence of ATP and the RR, the TCS complex is formed in the dark-adapted state of the signaling BphP (Fig. 2 a, movie S1). Compared to the cryo-EM density of the intact SaBphP2 (*13, 21*), further cryo-EM density is visible, corresponding to the RR, however, only on one side of the enzymatic domain (Fig. 2 a, b). For an interpretation of this density, the RR crystal structure (Fig. 2 c, fig. S1 b) was used. Two crystallographic RR_SaBphP2_ monomers forming a dimer can be placed into the cryo-EM density. Since the RR dimer binds to one side of the 4-helix bundle region of the BphP (Fig. 2 a,b, Fig. 3 a-c), both HK lobes reposition to allow for space for the binding of the RR (fig. S2 b). The resolution in the entire C-terminal region of the BphP, including HK domain (4.2 Å) is sufficient to unambiguously trace the amino acid backbone through the 4-helix bundle and through the HK lobes (Fig. 3 a, fig. S2 a, movies S1 and S2).

**Fig. 2.**
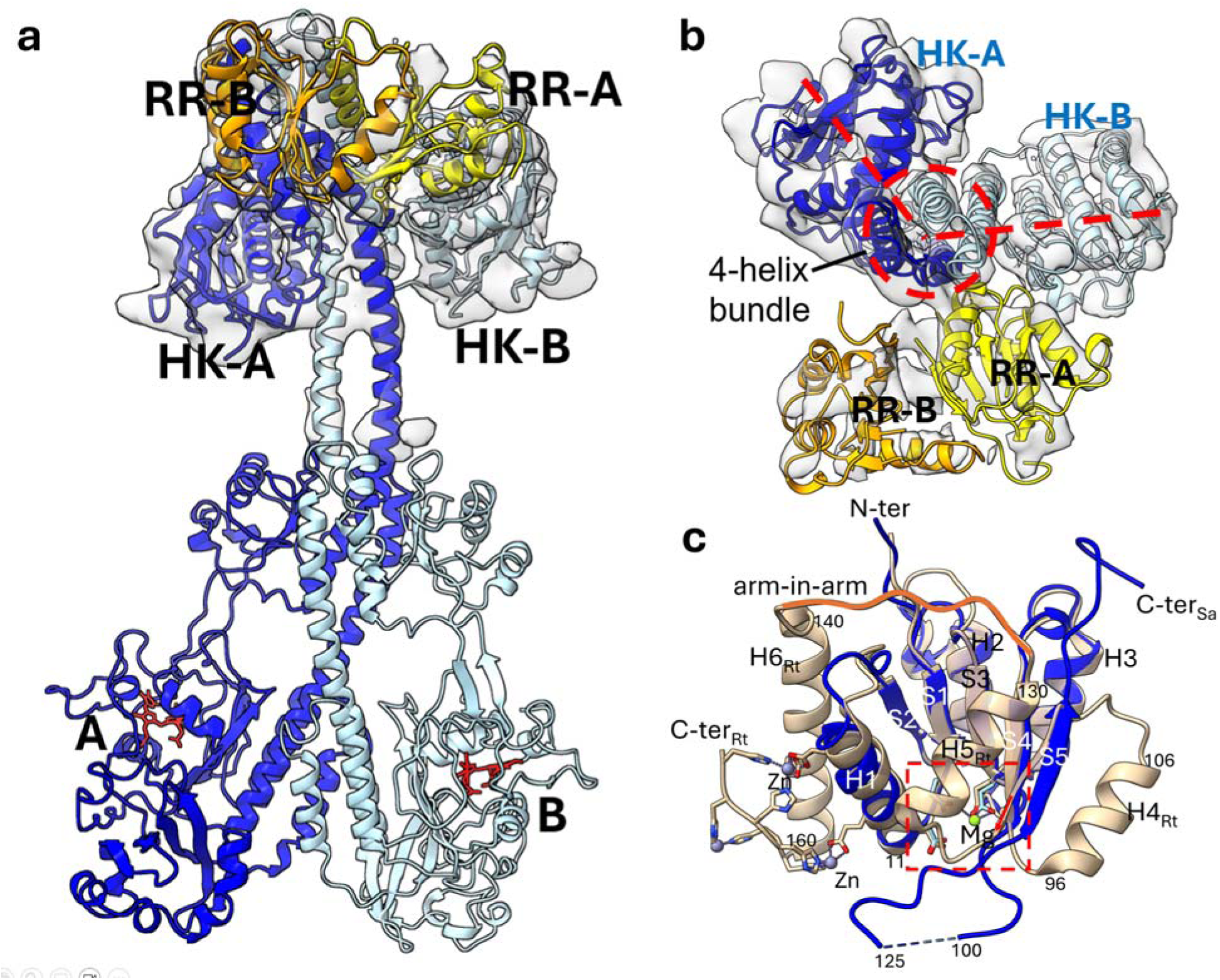
The SaBphP2-RR complex. **(a)** The RR_SaBphP2_ bound to the full length SaBphP2-Pr/Pr homodimer. Protomers A and B of SaBphP2 are blue and light blue, respectively. The BV chromophors are marked (red). The RR subunits A and B are shown in yellow and orange. **(b)** View from the top. The HK lobes are oriented at an angle (red dashed lines) to accommodate the RR-A which fits to a grove formed by the 4-helix bundle (marked) and HK-B. RR-B does not interact with the SaBphP2. **(c)** The crystal structure of the RR_SaBphP2_ (blue) overlayed on the *Ramlibacter tataouinensis (Rt)* BphP associated RR_RtBphP_ (light brown). The additional helices in RR_RtBphP_ and the arm-in-arm dimer interface (orange) can be easily identified, H4_Rt_ to H6_Rt_ are absent in RR_SaBphP2_. Red dashed box: the Asp rich region in RRs with magnesium^2+^ (Mg) bound. The N- and C-termini are marked for the two structures.

**Fig. 3.**
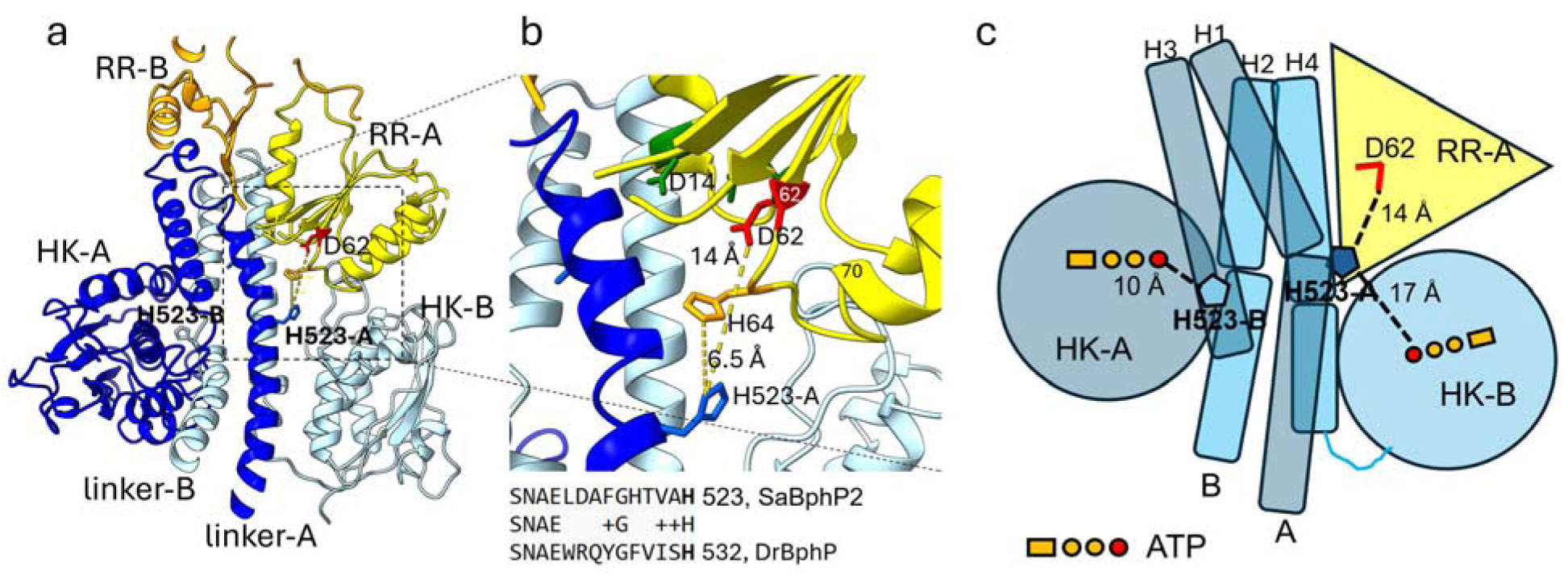
Details of RR binding to the SaBphP2 HK. **(a)** The SaBphP2 HK including the linker helices (linker A and linker B) and the HK lobes are shown, for protomer A, blue, and B, light blue, respectively. The bound RR is shown in yellow (subunit A) and orange (subunit B). The putative phosphorylation sites in the HK and the RR are marked (H523-A and D62, respectively). **(b)** The region within the dashed box is magnified. H523-A, and RR residues D14 and D62 are marked. The loop from amino acid residues 62 to 70 contains H64 that is located closer (6.5 Å) to the linker His523-A than the putative phosphate receptor D62 (14 Å). Bottom: A sequence alignment between SaBphP2 and DrPphP identifies the conserved H523 in SaBphP2. **(c)** Cartoon view derived from panel (a). Colors are similar to those in panel (a). Distances between the important residues His523 and the RR residue D62 are shown. Distances from the ATP molecules (γ-P_i_ in red) in the HK lobes to the respective H523 residues are also shown. Individual helices of the 4-helix bundle are numbered H1 to H4.

### SaBphP2 in the presence of ATP and the RR with light illumination

When ATP and the RR are added to SaBphP2 in the presence of ambient light, the TCS complex does not form– although the cryo-EM map (Fig. 4 c) shows that the signaling kinase has advanced through structural changes, previously characterized by absorption of a single photon in one or both protein monomers (*21*). The transition from the dark-adapted Pr/Pr homodimer to the light-triggered signaling Pr/Pfr heterodimer involves a ∼180° rotation of the entire HK domain (*13*). In the process, the linker helices partially unwind and the HK lobes interchange position (Fig. 1 a, Fig. 4 c). Absorption of a second photon results in an all-parallel linker helix configuration and formation of a fully light-adapted homodimer denoted as Pfr/Pfr (*21*). Like that of the Pr/Pfr heterodimer, the cryo-EM map of the Pfr homodimer (Fig. 4 d) lacks evidence for complex formation with the RR.

**Fig. 4.**
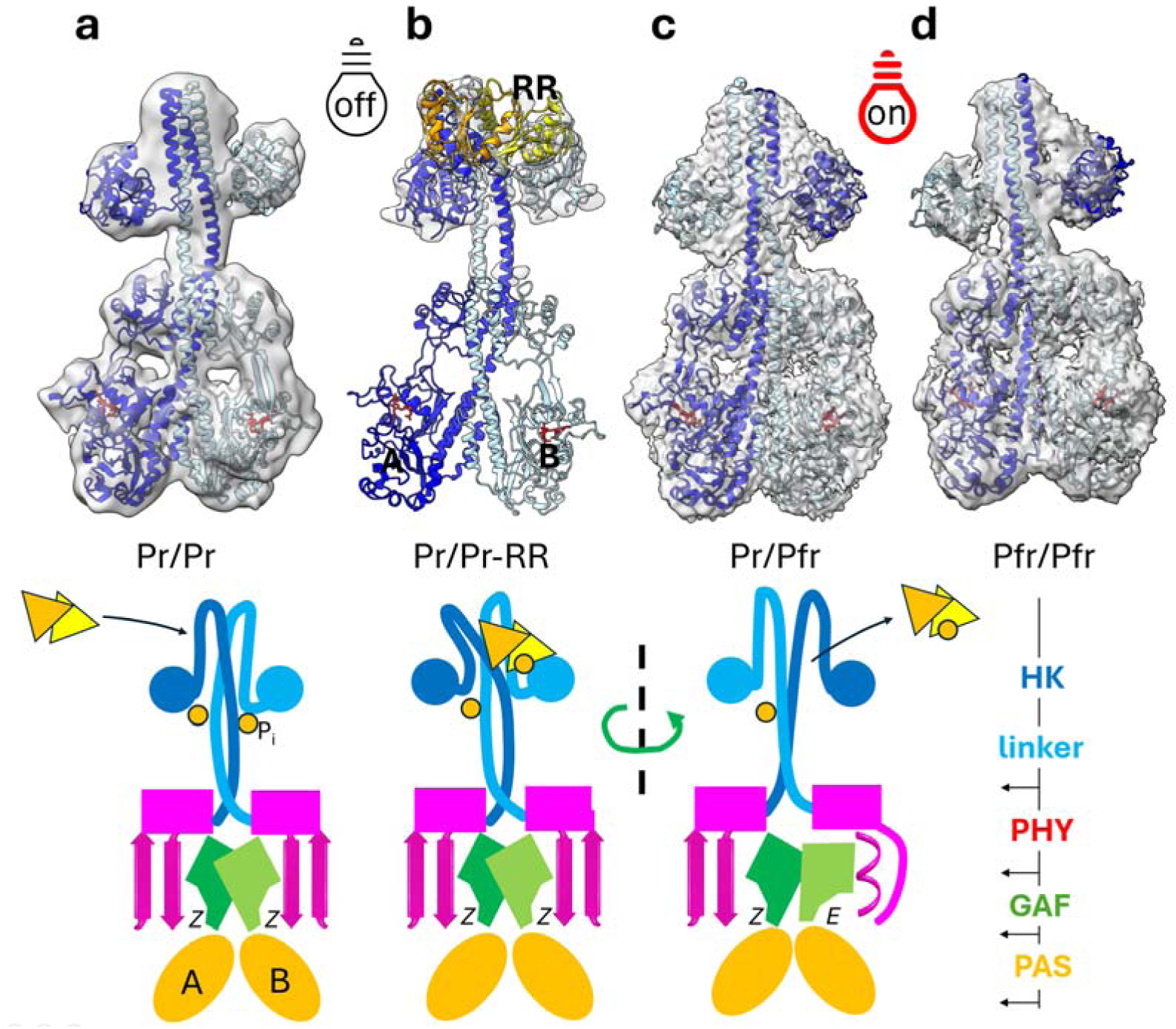
TCS signaling pathway consisting of SaBphP2 and its cognate RR in dark and light adapted states. **Upper panels**: cryo-EM structures for various photocycle states. Cryo-EM density displayed in transparent light gray (for (a), (c) and (d) at lower contour level). **(a)** The Pr/Pr homodimer (*13*) **(b),** the Pr/Pr-homodimer (protomer A in blue, protomer B in light blue) in complex with the RR (yellow and orange). **(c)** The Pr/Pfr heterodimer structure obtained from ambient light exposure in the presence of ATP and the RR, (d) the Pfr/Pfr homodimer structure after 660 nm light irradiation in the presence of ATP and the RR. No additional cryo-EM density for the RR can be observed in (c) and (d). **Lower panels:** Signaling pathway, derived from the structures of the upper panels taking into account the results from phosphotransfer assays. **(a)** BphP_Pr/Pr_ self-phosphorylates (circles marked by P_i_). The RR can bind. **(b)** The RR binds as a dimer, but only with one subunit. The binding induces changes in the HK structure that bends to the left. A phosphate is transferred to the RR. **(c)** Light absorption causes the isomerization of the BV to the *E*-configuration in one of the BphP protomers. This causes the β-sheet to α-helix transition of the sensory tongue, a large structural change of the BphP dimer interface, and an unwinding of the coiled-coil linker helices. The HK rotates by 180° (dashed line and green arrow). The Pr/Pfr heterodimer does not bind the RR. The domain composition is displayed on the right.

### The Crystal Structure of the RR_SaBphP2_

The RR_SaBphP2_ crystallizes (fig. S1) in spacegroup P4_3_2_1_2 with a single copy (a monomer) in the asymmetric unit (Fig. 2 c). The structure was solved by molecular replacement (MR). At a first glance, the architecture is similar to existing structures of BphP associated RRs (*22–25*) (Fig. 2 c) and other RRs with a canonical 5-stranded β-sheet (*26–28*). However, the RR_SaBphP2_ structure consists of only three α-helices and a five-stranded β-sheet (Fig. 2 c) where four of the strands are parallel but the fifth is antiparallel (fig. S1 c). Other BphP-associated RRs from *Ramlibacter tataouinensis* (RR_RtBphP_)(*29, 30*), the *Agrobacterium tumefaciens* (*23*) RR_AtBphP_ (*22, 29*) and the *Deinococcus radiodurans* RR_DrBphP_ (*31*) show a canonical 5-stranded β-sheet with all β-strands parallel (Fig. 2 c, fig. S1 c). The 5^th^ β-strand leads to helix H5 that is located on top of the β-sheet (Fig. 2 c). In some RRs, the c-terminal end of H5 is followed by a straight section of ∼10 AA residues that either ends in the C-terminus (RR_AtBphP_)(*29*) or extends to a long C-terminal helix H6 (RR_RtBphP_)(*29*) (fig. S1 c). The straight sections between H5 and H6 mutually forms an arm-in-arm dimer interface (*29*). Of the 6 helices found in RR_RtBphP_, only 3 analogous α-helices are present in RR_SaBphP2_ (Fig. 2 c, fig. S1 c). Interactions through the arm-and-arm interface do not occur in RR_SaBphP2_ dimers. Instead, the loop that leads to the 5^th^ β strand in one of the subunits makes contact with the 5^th^ β strand of the neighboring subunit which is related by crystallographic symmetry. This results in a rather large dimer interface from residue 129 to 138 (fig. S1c). Not surprisingly, when unpruned structures of various RRs with 5 or 6 helices were used as MR models, the MR failed due to unacceptable clashes. A crystallographic dimer can be obtained by reconstructing the content of the unit cell using the symmetry operations of the P4_3_2_1_2 space group. This dimer can be oriented as a rigid body into the cryo-EM density, the resulting structure is essentially identical to that obtained by placing individual monomers (fig. S1 b).

In the RR_SaBphP2_ a large loop from residues 100 to 125 remains structurally uncharacterized (Fig. 2 c, fig. S1 b,c). The loop protrudes into the large central pore left by the tetrameric arrangement in the unit cell (fig. S1 b). Zinc binding sites formed by the C-terminal helix and its extension in the RR_RtBphP_ do not exist in the RR_SaBphP2_. In RR_RtBphP_ and RR_AtBphP_ a magnesium (Mg^2+^) site is located at conserved Asp residues (Asp64 and Asp65 in the two structures, respectively). The equivalent site in the RR_SaBphP2_ structure is Asp62. Asp62 is part of an Asp rich structural motif found in all RRs. Asp62 is considered the phosphorylation site(*29*), in proximity to a conserved histidine (His523) located in the linker region, close to the HK lobes of SaBphP2 (Fig. 3 a).

### Kinase Activity Assay

Phosphotransfer from the HK to the RR(*2*) was examined under different dark/light conditions in order to capture SaBphP2 in the distinct signaling states, the dark-adapted Pr/Pr homodimer, the signaling Pr/Pfr heterodimer and the Pfr/Pfr homodimer. A phosphorylated RR band appears in PhosTag polyacrylamide gel electrophoresis gels immediately after mixing the RR with the dark-adapted SaBphP2 Pr/Pr homodimer (Fig. 5 a). The amount of phosphorylated RR (RR-P) decays with a lifetime of 1 min (inset in Fig. 5 a) which is similar to lifetimes reported for phosphorylated RRs such as the CheY and others(*2, 32–34*). The incubation mixtures contain elevated quantities of ATP (8 mM) to make sure that the SaBphP2 can effectively autophosphorylate and that the kinase activity continues after phosphate transfer to the RR. The appearance of a phosphorylated RR species at the earliest time point suggests that phosphotransfer occurs rapidly, faster than the decay of the RR-P. However, the self-phosphorylation rate of the BphP may be too slow to provide sufficient phosphate to compensate for the RR-P decay. When the experiment is repeated under ambient white-light conditions, which promote accumulation of the signaling Pr/Pfr heterodimer of the SaBphP2, phosphotransfer is no longer observed (Fig. 5b). This is corroborated by the cryo-EM density, which is free of RR occupancy (Fig. 4 c). Although the Pr/Pfr species autophosphorylates (*21*), the RR does not interact with the BphP sufficiently to receive a phosphate. Under direct red-light (660 nm) illumination conditions, when SaBphP2 is in the Pfr/Pfr homodimer state, the HK also does not transfer a phosphate (Fig. 5 b) and, in accordance with our cryo-EM experiments, it also does not bind the RR (Fig. 4 d).

**Fig. 5.**
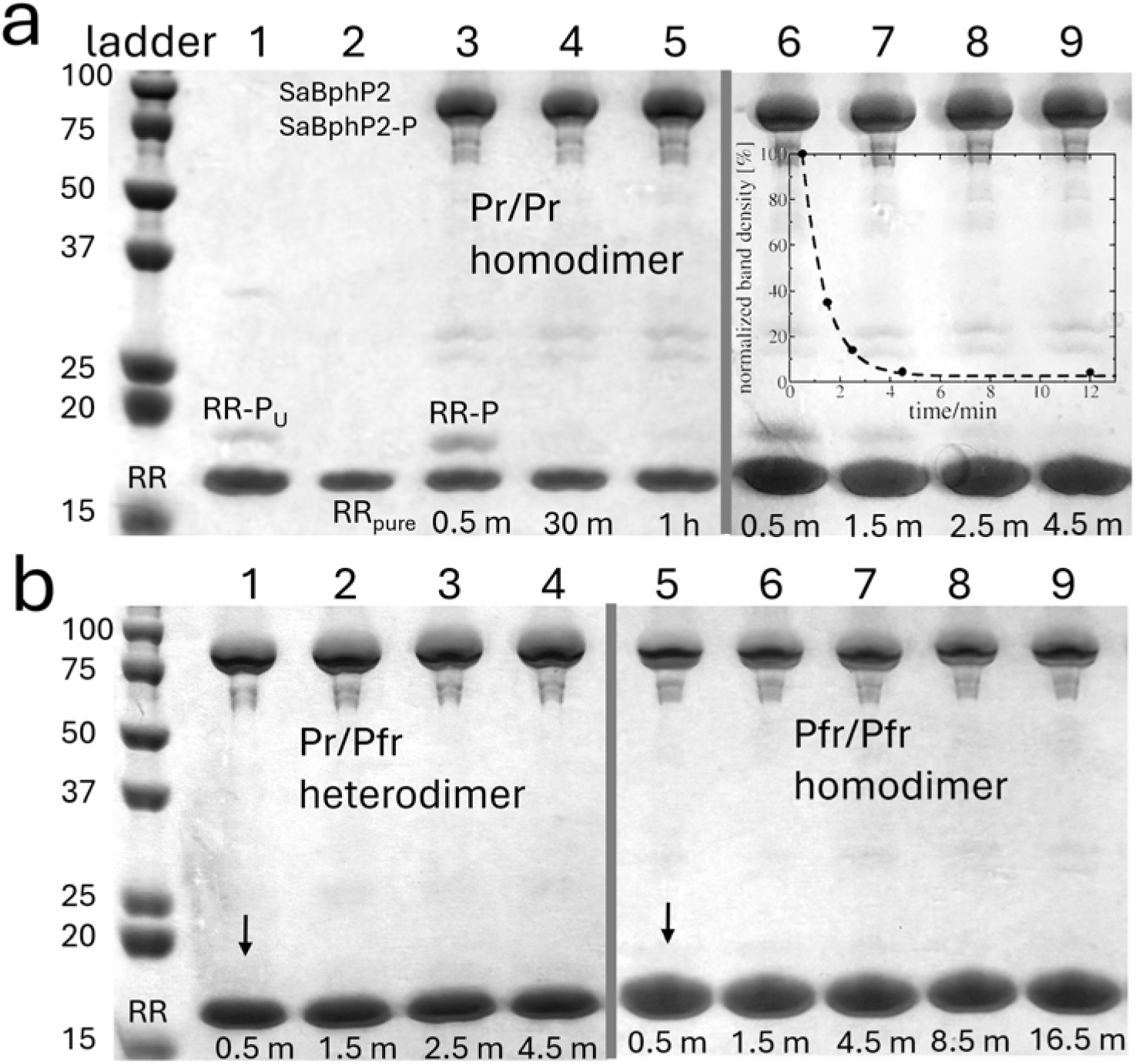
PhosTag polyacrylamide gel electrophoresis. The interaction of the RR with SaBphP2. **(a)** Interactions with the SaBphP2 Pr/Pr homodimer. Lane 1: unspecifically phosphorylated RR (RR-P_U_). Lane 2: FPLC purified RR (RR_pure_); lane 3: SaBphP2 + ATP + RR, 0.5 min after mixing; lane 4 and 5: 30 min and 1 h after mixing, respectively. The experiment is reproduced with samples drawn 0.5 min to 4.5 min after mixing (lanes 6 to 9, respectively). The phosphorylated RR (RR-P) band gradually vanishes with a lifetime of 1 min, see inset. **(b)** Interactions with the Pr/Pfr heterodimer, lanes 1 - 4, and the Pfr/Pfr homodimer lanes 5 - 9. Incubation times are marked for the various lanes. Transphosphorylation from SaBphP2 to the RR does not occur. The arrows denote the missing bands of the RR-P.

## Discussions

### Asymmetric RR association with the HK of SaBphP2

Our cryo-EM and HK activity assays confirm the formation of the TCS complex, composed of SaBphP2 and the RR, exclusively in the absence of the light signal. Movie S2 displays the cryo-EM density at lower contour level which shows the positions of the HK lobes, the structure of the linker helix and that of the 4-helix bundle in detail. The coiled-coil topology of the linker helices is a signature of the Pr/Pr state of SaBphP2. The HK-B lobe is displaced substantially as the RR binds (fig. S2 b). As a result, a binding interface around residue 700 is exposed that allows the RR dimer to bind into the cleft between the HK-B lobe and the 4-helix bundle helix #4 (H4) (Fig. 3 c). Only one monomer of the RR-dimer (RR-A) engages with the SaBphP2 HK. The loop in RR-A that spans from residue 62 to 68 interacts with a loop in the HK-B lobe of SaBphP2 around residue 700 (fig. S2 b). The N-terminal helix of the RR around Asp14 (Fig. 3 b) contacts the SaBphP2-A at the linker helix near Gln534-A and interacts with SaBphP2-B around residue 555 (located in H4). RR subunit B (RR-B) does not interact with the SaBphP2 HK. A small part of the density of the flexible loop can be identified in RR-B, but the loop seems to interact only with RR-A. The same flexible loop region in RR-A appears to wrap around the linker helix of SaBphP2-A. The exact locations of the loop regions, further away from the RR dimer, are difficult to determine, presumably due to structural disorder, which reflects an equivalent disorder in the protein crystal.

Both HK lobes, in particular the HK-B lobe (fig. S2 b), undergo substantial reorientation to accommodate RR binding, consistent with an induced-fit mechanism (movie S3). Sequence alignment of the SaBphP2 linker region with other BphPs (Fig. 4 b) indicates SaBphP2 His523 as the site for autophosphorylation. Further, it has been shown that the phosphate is transferred to an Asp residue in the RR where it is surrounded by Asp, Gln and Ser/Thr residues that also coordinate a Mg^2+^ (*27, 29, 32, 35*). By structural comparison, the important residues in the RR_SaBphP2_ are Asp62, Glu9, Asp14, Ser11 and Ser93 (Fig. 3 b). The Asp62 is the residue which is predicted to be phosphorylated (Fig. 2 c, Fig. 3 c). Consequently, to achieve transphosphorylation from the BphP to the RR, the RR must be oriented so that Asp62 is in the vicinity of an autophosphorylation site that can transfer the phosphate. In SaBphP2 a conserved histidine (His523) is this site which is located in the linker region close to the HK lobes (Fig. 3 a). Indeed, in the SaBphP2-RR complex the RR is bound in a way that the Asp-rich region in the RR points towards the His523. Note that a phosphate group cannot be identified at Asp62 in the cryo-EM density. Apart from challenges due to limited resolution, this is likely because the lifetime of the phosphorylated RR is substantially shorter than the time required to prepare and vitrify the sample for cryo-EM. The distance from SaBphP2 His523 to RR Asp62 cannot be determined exactly since the resolution is insufficient to display side chains, but it is on the order of 14 Å (Fig. 3 c) which would be too far for effective phosphate transfer. However, RR His64, is much closer to SaBphP2 His523 (∼ 6.5 Å) (Fig. 3 b) and could act as a phosphate shuttle to the neighboring Asp62. We, therefore, hypothesize that RR His64 is a transient, primary acceptor of the phosphate, that aids in the transfer to RR Asp62. We predict that substituting His64 with another amino acid (such as Alanine) will abolish phosphate transfer.

### ATP binding

Since ATP has been provided to the incubation mixture that led to the formation of the SaBphP2-RR complex, ATP should be present in the HK lobes. In order to identify a putative ATP binding site at low resolution, we overlayed the crystal structure of a truncated HK fragment from the *Rhodopseudomonas palustris* BphP2 (RpBphP2) which was crystallized in the presence of ATP (*36*) on the SaBphP2-RR complex. In the RpBphP2 crystals strong electron density determines the exact position of the ATP molecules. The structures of the RpBphP2 HK lobes are essentially identical to those in the SaBphP2 HK determined from our cryo-EM experiments. Accordingly, we placed ATP molecules into unexplained cryo-EM densities near the ATP positions identified in the HK_RbBphP2_. The ATP binding sites are shown in fig. S2 c,d and depicted schematically in Fig. 3 c. Notably, the γ-phosphate (γ-P_i_) of the ATP in the HK-A lobe is 10 Å distant from His523-B (Fig. 3 c). In contrast, the ATP in the HK-B lobe (to which the RR is binding) is 17 Å remote from His523-A (Fig. 3 c). When the ATP is placed into the HK-B lobe of the unperturbed, dark-adapted BphP, the distance of its γ-phosphate to His523-A is only 7.4 Å (fig. S2 b). Accordingly, we propose that in the Pr/Pr homodimer self-phosphorylation is catalyzed effectively by the HKs of both protomers (fig. S2 b). RR binding leads to the reorientation of one of the HK lobes. As a result, the contact of ATP with His523-A is disrupted and the phosphorylated residue His-523-A is exposed to the RR (Fig. 3 c, fig. S2 b). Subsequently, the RR accepts the P_i_ from His523-A.

### Mechanism of TCS signaling

Our cryo-EM structures show that autophosphorylation must occur across the dimer interface, since, due to the twisted coiled coil helix geometry in the Pr/Pr homodimer, the HK-B lobe is in proximity to the linker helix of protomer A (linker-A, Fig. 3 a, Fig. 4 a). In turn, since HK-A is in the vicinity of linker-B, it can phosphorylate linker-B as well. This way, the across-protomer (in-*trans*) autophosphorylation established through site-directed mutagenesis in other BphPs (*36*) can naturally be understood. This result is further supported by autophosphorylation of EnvZ (*37*) and NtrB (*38*) in *trans*, that is the CA domain of one subunit provides the ATP and catalyzes the phosphorylation of the phosphoreceptor His of the other subunit. Our results are in contrast to conclusions of previously published, incomplete structures of TCS complexes (*8–10*) containing the cytosolic portion of the sensory HK and the corresponding RR. In these cases, the autophosphorylation reaction was proposed to occur in *cis* (*8*) where ATP bound to one subunit phosphorylates the His residue of the same subunit.

The TCS complex from *A. aurantiaca* as determined by cryo-EM, in combination with the results from PhosTag-PAGE, provides a molecular mechanism for light-controlled signal transduction mediated by SaBphP2 (movie S3). Based on the unperturbed structure of the SaBphP2 Pr/Pr homodimer (*13*), the His523 residues in both protomers are phosphorylated by the HK. We hypothesize that phosphorylation is necessary to attract the RR. When the RR binds, one of the HK lobes (HK-B) will reorient providing access for the RR to the phosphorylated His523-A. In the RR, a loop that includes the putative phosphorylation site, Asp62, comes into the vicinity of the His523-A (Fig. 3, Fig. 4 a,b). A phosphate can be transferred from the BphP to the RR. The phosphate is likely shuttled through RR His64 before it is transferred to Asp62 (Fig. 3 b). The resulting phosphate-monoester is only transiently stabilized by closeby RR residues.

The phospho-aspartyl is fragile (*33*) and decomposes (hydrolyses) within 1 min as evidenced from the corresponding PhosTag gels (Fig. 5 a). Since the linker-B helix is far from the RR binding site (Fig. 3 c), it does not interact with the RR residues His64 and Asp62. As the result, the RR cannot accept a phosphate from His523-B. Accordingly, the phosphotransfer is asymmetric: the phosphate is transferred to only one RR subunit at a time by only one protomer of the BphP.

In the presence of weak ambient light, the TCS complex is not formed. Red light absorption leads to the formation of the BphP Pr/Pfr heterodimer with a concomitant 180° rotation of the HK switching the HK lobes from one side of the BphP to the other. As a result, the Pr/Pfr heterodimer and also the Pfr/Pfr homodimer are unable to bind the RR (Fig. 4 c,d, Fig. 5 b). A once formed SaBphP2-RR complex likely dissolves with light activation (movie S3). The Pr/Pr homodimer is the only species that is involved in phosphate transfer to the RR (Fig. 5 a). The(quick) dephosphorylation of the RR seems to be an important way to prevent signaling in the absence of red light. So far, it remains an open question (*2, 34*) whether transient (1 min) phosphorylation of a single RR subunit within the dimer can generate a sufficiently strong and sustained signal to initiate a cellular response. It is also not known whether the RR remains a dimer or monomerizes upon dissociation from the SaBphP2. However, the structural similarity with CheY(*32, 35*) and other RRs that affect motility(*3*) suggest that RR_SaBphP2_ is involved primarily in the light-control of *A. aurantiaca* motility that results in swarming and fruiting body formation (*15*) (Fig. 6) as also recently demonstrated for a related myxobacterium, *Myxococcus macrosporus,* that contains a single BphP (*39*). This way, our results establish a direct link from early events in BphPs induced by light (*40, 41*) to light-driven signal transduction (*21*) and physiological responses.

**Fig. 6.**
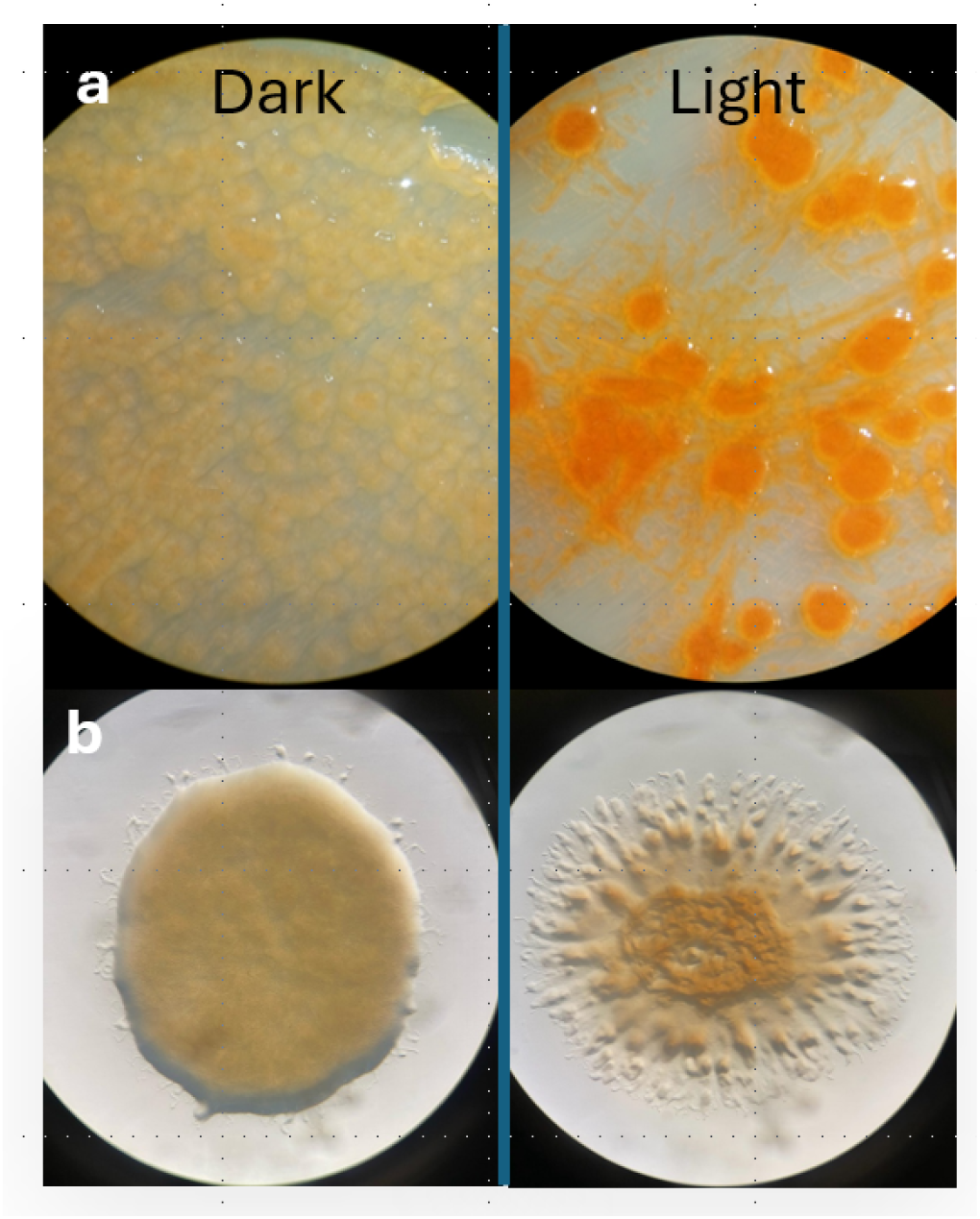
Photomorphogenesis in *A. aurantiaca*. Vegetative cells after 14 days (a) or 4 days (b) of cultivation on nutrient agar plates in constant dark vs light conditions as imaged by light microscopy. **(a)** Fruiting body formation and carotenoid synthesis occur only with light exposure. **(b)** Swarming is induced by light. In the dark cells remain sessile.

### Impact

Our results contribute to the understanding of HK–RR recognition in myxobacteria, specifically in light-signaling kinases. The BphP and the RR are constitutively expressed in *M. macrosporus* and likely in *A. aurantiaca*, regardless of the light condition (*39*). Therefore, when the bacterial cells are in the dark, BphPs and RRs are mostly associated, while in the presence of light, the BphP-RR complex disintegrates, and the BphP Pr/Pfr heterodimer remains the abundant species. Here, a molecular basis is provided for predicting and manipulating signaling specificity as proposed for other TCS systems (*5*), including the rational rewiring of said pathways (*42*). Although *A. aurantiaca* is non-pathogenic, the interactions between its HK and cognate RR provide fundamental insights into how TCS specificity and partner selection may govern downstream cellular behaviors in pathogens, including motility and biofilm formation (*43*) and can inform strategies for targeting analogous signaling pathways across various bacterial kingdoms (*6*). Furthermore, *A. aurantiaca* and most myxobacteria are predatory soil bacteria. Our results provide strong evidence that light control might be critical for biodiversity and health of the soil biome (*44*).

## Acknowledgments

We thank Niko McCracken and Dan Sierks (UWM) for help with the UWM HPC cluster.

## Author contributions

Conceptualization: MS, EAS

Methodology: CH, JM, WB, PK, MS, PE, DM, SK, SR, RA, DM

Investigation: JM, CH, WB, PE, PK Visualization: PK, MS

Funding acquisition: MS, EAS Project administration: MS, EAS Supervision: MS, EAS

Writing – original draft: MS, PK, EAS

Writing – review & editing: MS, EAS, PK and all other authors

## Data, code, and materials availability

The structure of the *Sa*BphP2–RR complex is available in the Protein Data Bank (PDB) (*45*) under accession code 39AA. The corresponding cryo-EM density maps are available in the Electron Microscopy Data Bank (EMDB) (*46*): the map reconstructed by local refinement of the PCM region under EMD-79261, the HK–RR region under EMD-79262, and the composite map combining the PCM and HK–RR regions under EMD-79271. The cryo-EM density map of the Pr/Pfr heterodimer in the presence of ATP and RR is available under EMD-79295, with the corresponding structure archived in the PDB with entry 39DH. The Pfr/Pfr homodimer map, determined in the presence of ATP and RR, is available under EMD-79296, with the corresponding structure in the PDB with entry 39BS. The structure and the structure factors of the *Sa*BphP2 response regulator are available in the PDB with entry 39AV. The *Sa*BphP2 and RR overexpression systems are available upon request.

## Supplementary Materials

Materials and Methods Figs. S1 to S4

Tables S1 to S2 References 47 -52

## Other Supplementary Materials for this manuscript include the following

Supplementary movies will be available once the manuscript is published in a peer reviewed journal

Movies S1 to S3

## Materials and Methods

### SaBphP2 and RR expression and purification

*Stigmatella aurantiaca* DW 4/3-1 was very recently renamed to *Allostigmatella aurantiaca* DW 4/3-1 (*47*) . In this manuscript and for the purpose of avoiding confusion with previous literature we refer to the *A. aurantiaca* bacteriophytochrome 2 as SaBphP2 rather than AaBphP2. The SaBphP2 and its cognate response regulator (RR_SaBphP2_) are both located in the same operon. Full-length (intact) SaBphP2 was cloned earlier and purified using a published protocol (*12, 21*) and stored at -80 °C in stabilization buffer (Tris/HCl 25 mM, 75 mM NaCl, pH 8.0). The gene of the RR was synthesized by GenScript with an N-terminal 6-His tag and a TEV protease cutting site, inserted into a pET30a expression system, and overexpressed in 5 L terrific broth at 22 °C in E. coli BL21-STAR (DE3).

After the overexpression of the RR, the cells were centrifuged at 5000 g for 10min at 4 °C and resuspended in 150mM NaCl, 20mMTris-HCl pH 8.0 with a protease inhibitor cocktail (Roche). The cells were lysed using sonication and centrifuged at 23 000 g for 1 h at 4 °C. The supernatant solution was added to Ni-NTA metal ion affinity chromatography columns (Takara Clontech). The protein was eluted with 200mM imidazole, 20mM NaCl, and 20mM Tris-HCl pH 8.0. After buffer exchange to 20 mM Tris, 150 mM NaCl, 1 mM EDTA, 1 mM DTT, pH 7.8, about 200 mg of RR (MW 17,442 Da) was obtained which was concentrated to 50 mg/ml and stored at -80 °C. A small fraction of the RR appeared phosphorylated during expression (Fig. 5 a, lane 1). Presumably, *E. coli* must have a kinase that was active with the RR_SaBphP2_ during overexpression. To separate the contamination, the RR was purified by anion chromatography using Q-sepharose at pH 8 using an Akta Prime FPLC chromatography system (Cytiva) featuring a standard gradient between 100 mM NaCl and 1 M NaCl (in 30 mM Tris/HCl, 0.5 mM DTT, pH 8.0). A preparation free of contamination (Fig. 5 a, lane 2) was obtained and concentrated to 15 mg/ml.

### Phosphorylation experiments using PhosTag polyacrylamide gel electrophoresis

Transphosphorylation of the RR_SaBphP2_ by the SaBphP2 Pr/Pr homodimer, the Pr/Pfr heterodimer and the Pfr/Pfr homodimer was investigated using sodium dodecyl sulfate (SDS) polyacrylamide gel electrophoresis containing PhosTag resin (PhosTag-PAGE). Pre-prepared PhosTag gels (12.5 % polyacrylamide, 50 μM PhosTag) were acquired from Fuji Film. The gels were run with 80 V at room temperature using a mini electrophoresis chamber (BioRAD). After completion, the gels were stained with Coomassie blue (BioRAD) and destained with a mixture of water, methanol (20%) and glacial acidic acid (20%). The gels were photographed using a iPhone camera (Apple). The density values of the bands containing phosphorylated RR (RR-P) were integrated by Claude (Anthropic). Sample preparation details are as follows: Stock solutions for MgCl_2_ (200 mM), MnCl_2_ (200 mM) KCl (250 mM) and ATP (50 mM) were prepared in stabilization buffer (Tris/HCl, pH 8, 100 mM NaCl, 0.5 mM DTT) and the pH adjusted with NaOH if needed. In an incubation volume of 40 μl, 1.5 μl of freshly thawed SaBphP2 (50 mg/ml) was mixed with MgCl_2_, MnCl_2_ and KCl stock solutions to final concentrations of 20 mM, 2 mM and 20 mM, respectively. To obtain a preparation of pure Pr/Pr homodimers, SaBphP2 was illuminated for 1.5 min by a 740 nm light diode (Thorlabs). After the illumination, ATP was added to a final concentration of 8 mM. The mixture was kept at room temperature (and in the dark) for about 10 min to allow for efficient autophosphorylation. After this, 7 μl of FPLC purified RR (15 mg/ml stock solution in stabilization buffer) was added. The mixture was illuminated by 740 nm light for an additional 0.5 min and stored on ice. 10 μl was immediately removed, mixed with sample buffer (Bio-Rad) at room temperature and stored on ice. To investigate the interaction of the Pr/Pfr heterodimer with the RR, all steps were conducted without pre-illumination of the *Sa*BphP2 and without protection from laboratory light. The interaction of the Pfr/Pfr homodimer with the RR was investigated using the same protocol as for the Pr/Pr homodimer, except that the mixture was illuminated with 660 nm light. The collected samples (15 μl) were applied to the PhosTag-gels. Results are shown for the various species (the SaBphP2 Pr/Pr homodimer, Pr/Pfr heterodimer and Pfr/Pfr homodimer) in Fig. 5. The unspecifically phosphorylated RR (Fig. 5 a) served as a convenient control.

### Preparation of SaBphP2 species in the presence of the RR for cryo-EM experiments

Purified SaBphP2 and the RR_SaBphP2_ were sent to New York Structural Biology center. Mixtures containing BphP, Adenosine Triphosphate (ATP) and RR were prepared in identical ways for 3 BphP species, the Pr/Pr homodimer, the Pr/Pfr heterodimer and the Pfr/Pfr homodimer. In a total volume of 50 μl, SaBphP2 (0.3 mg/ml) in stabilization buffer (Tris/HCl 30 mM, 50 mM NaCl, pH 8) was mixed with MgCl_2_ (20 mM), KCl (50 mM) and ATP (3 mM). To obtain the Pr/Pr homodimer, the mixture was illuminated for 2 min with 740 nm light from a 15 mW higher power LED (Thorlabs). After this, the RR (0.1 mg/ml) was added. All concentrations are final concentrations. The proteins, the salts and the ATP were obtained from stock solutions (2 mg/ml, 1.5 mg/ml, 50 mM, 250 mM, 500 mM for the SaBphP2, RR, ATP, MgCl_2_ and KCl, respectively). The Pr/Pfr heterodimer was prepared by thawing SaBphP2 from storage (-80 ^0^C) and exposing it to ambient laboratory light throughout the incubation. Particles corresponding to the Pfr/Pfr homodimer were detected in the same preparation and separated by classification (see below). 3 μl of the mixtures were applied to holey grids, blotted and plunge frozen using a Vitrobot. Preparations of the Pr/Pr and the Pfr/Pfr homodimers were handled under green (500 nm) safety light.

### Cryo-EM

Cryo-EM movies were collected with a Thermo Fisher Scientific Titan Krios electron microscope as previously reported (*12, 21*). The movies were sent back to UWM for further processing using CryoSPARC v. 4.7.1(*48*). Data collection statistic including the obtained resolution limit is reported in table S1.

### The SaBphP_Pr/Pr_-RR complex

A flowchart illustrating the cryo-EM data-processing workflow is shown in fig. S3. A total of 14,709 movies were imported into cryoSPARC v4.7.1, followed by motion correction and contrast transfer function (CTF) estimation. Micrographs were filtered based on CTF fit resolution and relative ice thickness, resulting in 8,473 micrographs retained for downstream processing. Initial particle picking was performed using blob picker with a diameter range of 70– 250 Å, yielding 3,303,145 candidate particles. Iterative 2D classification was used to remove poorly resolved particle classes, retaining 141,776 particles. The resulting 2D class averages were used as templates for template-based particle picking. Subsequent iterative 2D classifications and particle selection resulted in 243,514 particles. These particles were subjected to 3-class ab initio reconstruction followed by heterogeneous refinement, producing an initial reconstruction at 5.15 Å resolution from 80,176 particles. Subsequently, neural network-based particle picking using Topaz was performed, with the model trained on the template-picked particle set, resulting in 166,268 particles. Nonuniform refinement using the 5.15 Å reconstruction as an initial reference yielded a map at 6.6 Å resolution. The particles were then subjected to 3D classification into two classes, and the lower-quality class containing 45.1% of the particles was discarded. Nonuniform refinement of the retained particles improved the reconstruction to 4.71 Å resolution. Subsequent reference-based motion correction followed by nonuniform refinement further improved the global resolution to 4.51 Å. To improve density quality and local resolution, focused refinement was performed on both the PCM region as well as the HK-RR region. Masks for both regions were generated in UCSF ChimeraX, imported into cryoSPARC, and used for particle subtraction followed by local refinement. These focused refinements yielded maps with resolutions of 4.29 Å and 4.37 Å, respectively.

### The Pr/Pfr and Pfr/Pfr species of SaBphP2 in the presence of ATP and the RR

A flowchart illustrating the cryo-EM data-processing workflow is shown in fig. S4. A total of 22,484 movies were collected and imported into cryoSPARC, where they were motion-corrected and CTF-estimated. Following manual curation, 18,324 micrographs of sufficient quality were retained for downstream processing, and the remainder were discarded. Initial particle picking was performed using the blob picker, yielding 5,372,341 particles. Iterative rounds of 2D classification and class selection reduced this set to 185,641 particles. These particles were subjected to 2-class ab-initio reconstruction followed by heterogeneous refinement. Non-uniform refinement of class 0 produced a 3.90 Å map, which was used to generate templates for template-based particle picking. This second round of picking yielded 5,396,553 particles, which were reduced to 457,375 particles through iterative 2D classification and selection. These particles were subjected to 3-class ab-initio reconstruction. Non-uniform refinement of class 0 yielded a 3.66 Å Pr/Pfr map, which was in turn used to generate a new set of templates. A subsequent round of template-based picking produced 660,169 particles after iterative 2D classification. This particle set was subjected to a second round of 3-class ab-initio reconstruction, and classes 1 and 2 were further processed by heterogeneous refinement. Non-uniform refinement of the resulting volumes produced a 3.68 Å Pfr/Pfr–PCM map with well-resolved PCM density and a 3.56 Å Pr/Pfr map. Reference-based motion correction and local/global CTF refinement, followed by a final round of non-uniform refinement, improved these maps to 3.57 Å (Pfr/Pfr, 255,021 particles) and 3.46 Å (Pr/Pfr, 244,748 particles), respectively. The 3.46 Å Pr/Pfr map was of sufficient quality to support model building.

While the Pr/Pfr map showed clearly resolved density for the histidine kinase (HK) region, the Pfr/Pfr map — despite well-resolved PCM density — lacked interpretable HK density. To improve this region, a new round of template-based picking was performed using particles originally identified by the blob picker as templates. This yielded 9,683,224 particles, of which 5 million were extracted; iterative 2D classification and selection reduced this to 1,629,295 particles. This particle set was subjected to 3-class ab-initio reconstruction, followed by heterogeneous refinement of classes 0 and 1. Class 0 corresponded to Pr/Pfr, with poorly resolved HK density, whereas class 1 corresponded to Pfr/Pfr with clearly defined HK density. Non-uniform refinement of class 1 yielded a 3.59 Å Pfr/Pfr map from 895,936 particles. Although one HK lobe was well resolved in this map, the second lobe remained poorly defined at higher contour levels. To remove residual junk particles, three junk volumes were generated and used as decoy classes in a subsequent heterogeneous refinement job. Following this heterogeneous refinement and a final round of non-uniform refinement, a 3.74 Å Pfr/Pfr map, using 188,221 particles, with improved density for both HK lobes was obtained. This map was used for atomic model building of the Pfr/Pfr state.

### Cryo-EM Structure determination and structural refinement

To determine the structure of the *Sa*BphP2_Pr/Pr_-RR complex, the structural model of the Pr/Pr homodimer determined by Karki(*13*) was used as initial model. Caused by the RR binding, this model required extensive modifications within the HK region to fit the density. Additional cryo-EM density indicated the presence of the RR. Two crystallographic monomers were independently oriented into the density using ChimeraX(*49*). An essentially identical result was obtained when a crystallographic dimer was placed as a rigid body into the density. The complete model was refined against the density map using Isolde (*50*) and Phenix v1.21.2 (*50*) (table S1).

The Pr/Pfr heterodimer determined by Malla et al. (*21*) fits the Pr/Pfr cryo-EM density well. No additional density that would indicate RR binding can be identified. The cryo-EM density of the Pfr/Pfr homodimer is contingent in the HK region. The partial model of Pfr/Pfr homodimer determined previously (*21*) is placed in the density. The partial model is completed by adding the structures of the missing linker helices and the HK lobes. Because the cryo-EM density obtained here is of sufficient quality to model the previously unresolved regions, the missing regions were taken from the previously determined Pr/Pfr structure (*13*) and manually positioned into the Pfr/Pfr cryo-EM density using ChimeraX. The missing regions were then connected to the existing model by creating the appropriate covalent bonds, followed by real-space refinement in Phenix against the cryo-EM density. The cryo-EM densities of the Pr/Pfr and Pfr/Pfr species are inspected at lower contour levels. Additional density that might indicate the presence of the RR is not observed (Fig. 4). Movie S3 depicts a morph between four species consisting of the Pr homodimer, the Pr/Pr-RR complex, the Pr/Pfr heterodimer and the Pfr homodimer to outline the structural transitions more clearly. Data statistics are shown in table S1.

### Crystallization and structure determination of the RR_SaBphP2_

The RR_SaBphP2_ was crystallized by mixing an equal amount of Hampton Crystal Screen Cryo #18 (0.16 M magnesium acetate, 0.08 M Na cacodylate trihydrate, pH 6.5, 16 % PEG 8000 and 20 % glycerol) with an equal amount of the RR (50 mg/ml) in a sitting drop vapor diffusion experiment. Large crystals (100 μm, fig. S1) grow over the course of a few days. To determine a high-resolution structure of the RR, the crystals were immersed in the Hampton Crystal Screen Cryo #18 buffer and sent in Eppendorf tubes to the European Synchrotron Radiation Facility (ESRF) beamline ID-23-2. Diffraction patterns were collected from one crystal kept at T=100 K using 14.2 keV (0.87 Å) X-ray radiation. The patterns were analyzed by Imosflm (v7.4.0)(*51*). Structure factor amplitudes were obtained up to a resolution of 2.3 Å. The spacegroup was P4_3_ 2_1_ 2 with unit cell parameters a = b = 86.15 Å and c = 39.04 Å. The data collection statistics is shown in table S2. Molecular replacement (MR) was attempted using a variety of published RR structures as well as with a model prepared by AlphaFold3 (*52*). All models did not pack unless some of the peripheral helices were removed. Otherwise, severe clashes with symmetric equivalent RR molecules prevented a molecular replacement solution. The fifth β-strand (of the canonical 5-stranded β-sheet in the RRs) was also removed from the search models. After a successful molecular replacement using phaser-MR implemented in phenix v.1.21-5207 (*50*), strong electron density of the fifth β-strand is present from which amino acid side chains can be identified. The succession of the amino acids is oriented in the opposite direction compared to that observed for the analogous β-strand in other RRs and it is also antiparallel to that predicted in the AlphaFold3 model. Helices H4 to H6 present in some RR structures are all missing. Hence, dimer-interface interactions previously observed in other RRs (*26–28*) and for BphP associated RRs (*29*) are not observed. Instead, between β-strands 4 and 5 of the five membered β-sheet a stretch of 43 AA residues forms a partially disordered loop (fig. S1 c). A portion of this loop together with β-strand 5 are part of mutual interactions with a crystallographically symmetric molecule. This crystallographic dimer can also be identified and positioned in the cryo-EM density (fig. S1 b). The positions of ∼25 AA residues in this loop could not be traced faithfully and are omitted from the model. The incomplete model was refined in Phenix to R_cryst_/R_free_ of 22%/27% (table S2). When the 5 – helix AlphaFold3 predicted RR model is positioned in the cryo-EM map, density for helices H4 and H5 (Fig. 2 c) is absent confirming the validity of the crystal structure determined here.

### Vegetative cultures of *A. aurantiaca*

*A. aurantiaca* was streaked out from glycerol stocks onto nutrient-rich bacto agar plates containing 1% Tryptone, 0.2% MgSO_4_.7H_2_O, Bacto Tryptone, prepared in HPLC-grade water and incubated at 30 °C for 5 days. The components were mixed in HPLC grade water, and the pH was adjusted to 7.2 before adding agar. If the pH exceeded 7.2, it was corrected using NaOH. The medium was then autoclaved for 15 minutes. For liquid cultures (Tryptone broth), the same ingredients were used and the same procedure was followed, except that the agar was omitted. Liquid cultures were incubated at 30°C and shaken at 170 rpm until they reached the desired optical density (OD_600_). Samples from both agar plates and broth cultures were Gram-stained and examined with a light microscope to assess cell structure. Following this, cultures were centrifuged at 12,000rpm for 15 minutes at 4 °C. Cell pellets were resuspended in fresh media, and 5uL of each suspension was spotted onto separate quadrants of a divided agar plate. The spotted plates were incubated at 30°C and observed for 14 days under two environmental conditions: constant darkness (DD) and constant white LED light (LL). Bacterial growth and development were monitored using a Zeiss microscope and images were captured with the Zeiss Zen software.

## Supplementary Figures

**Fig. S1.**
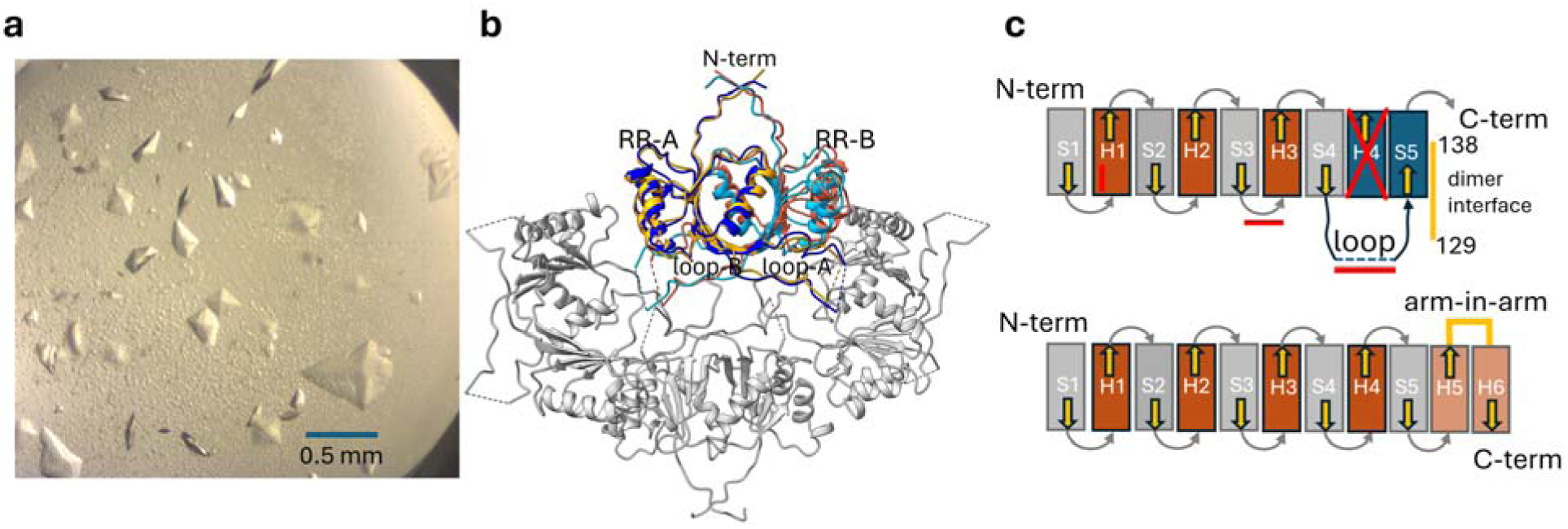
The RR_SaBphP2_ crystal structure. **(a)** Bi-pyramidally shaped crystals of RR_SaBphP2_ in a sitting drop reflecting the tetragonal symmetry. **(b)** Unit cell content by applying symmetry operations: the RR forms a crystallographic dimer (monomers are related by a symmetry operation) which is shown in blue and light blue. The RR dimer identified by cryo-EM (yellow, orange) is overlayed for comparison. The structures and the dimer interfaces of the two dimers are essentially identical. **(c)** Comparison of the structural topology of the RR _SaBphP2_ (upper) to that of the RR_RtBphP_ (lower). Arrows denote increasing sequence numbers starting from the N-terminus (N-term), S1-S5: strands of the 5-membered β-sheet, H1-H6: helices in the different RRs. Red bars: interactions with the BphP, yellow bars: dimer interfaces (for the RR_SaBphP2_ sequence numbers for the interface are provided).

**Fig. S2.**
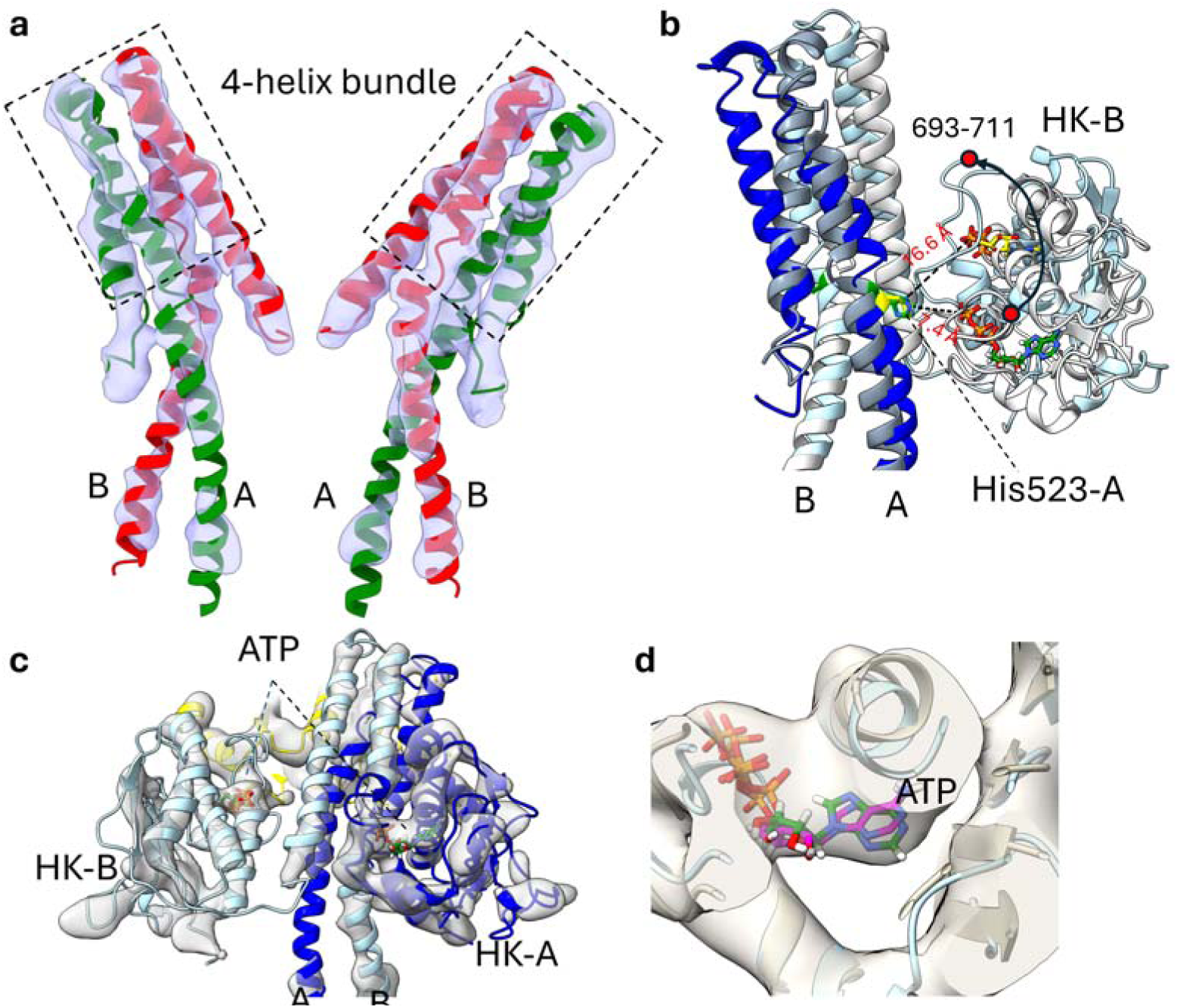
Structural details. **(a)** The linker and 4-helix bundle region in the SaBphP2-RR complex from the front and the back, respectively. SaBphP2 protomers are marked A and B. The HK lobes and the RR are removed. Cryo-EM density is shown in light blue. The cryo-EM density unambiguously shows the structure of the 4-helix bundle and the loops that lead to the HK lobes, respectively. **(b)** Structural displacement of the HK-B lobe in the SaBphP2-RR complex (blue, the RR is not shown) compared to the unperturbed, dark adapted SaBphP2 (silver). The red dot denotes the large loop that provides a binding interface for the RR after a large structural reorientation of the lobe (arrow). The position of ATP is shown before and after the structural change. Distances to the γ-phosphates are marked. They are 7.4 Å in the unperturbed structure and 16.6 Å in the SaBphP2-RR complex, respectively. **(c)** The position of the ATP in the SaBphP2 HK lobes. The structure is viewed from the backside to better identify the ATP position. The RR (yellow) is located behind HK-B. **(d)** Cryo-EM density (transparent grey) of the HK ATP binding moiety (blue structure) in protomer B. An ATP molecule (pink) fits the density. In comparison, the ATP molecule identified by X-ray crystallography in the RpBphP2 HK lobe (grey) is shown in green.

**Fig. S3.**
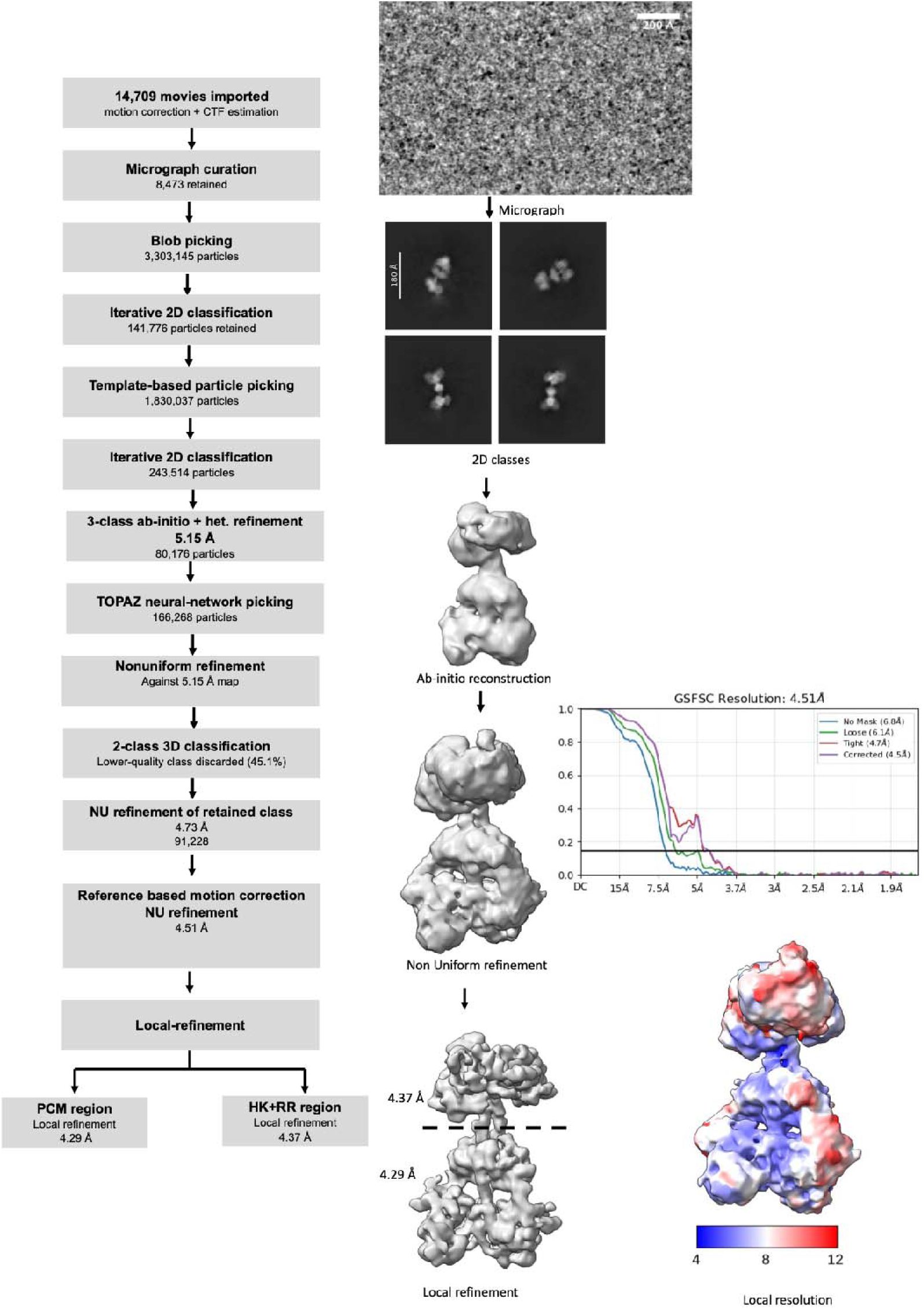
Flow chart for the processing of Cryo-EM data collected on the SaBphP_Pr/Pr_-RR complex.

**Fig. S4.**
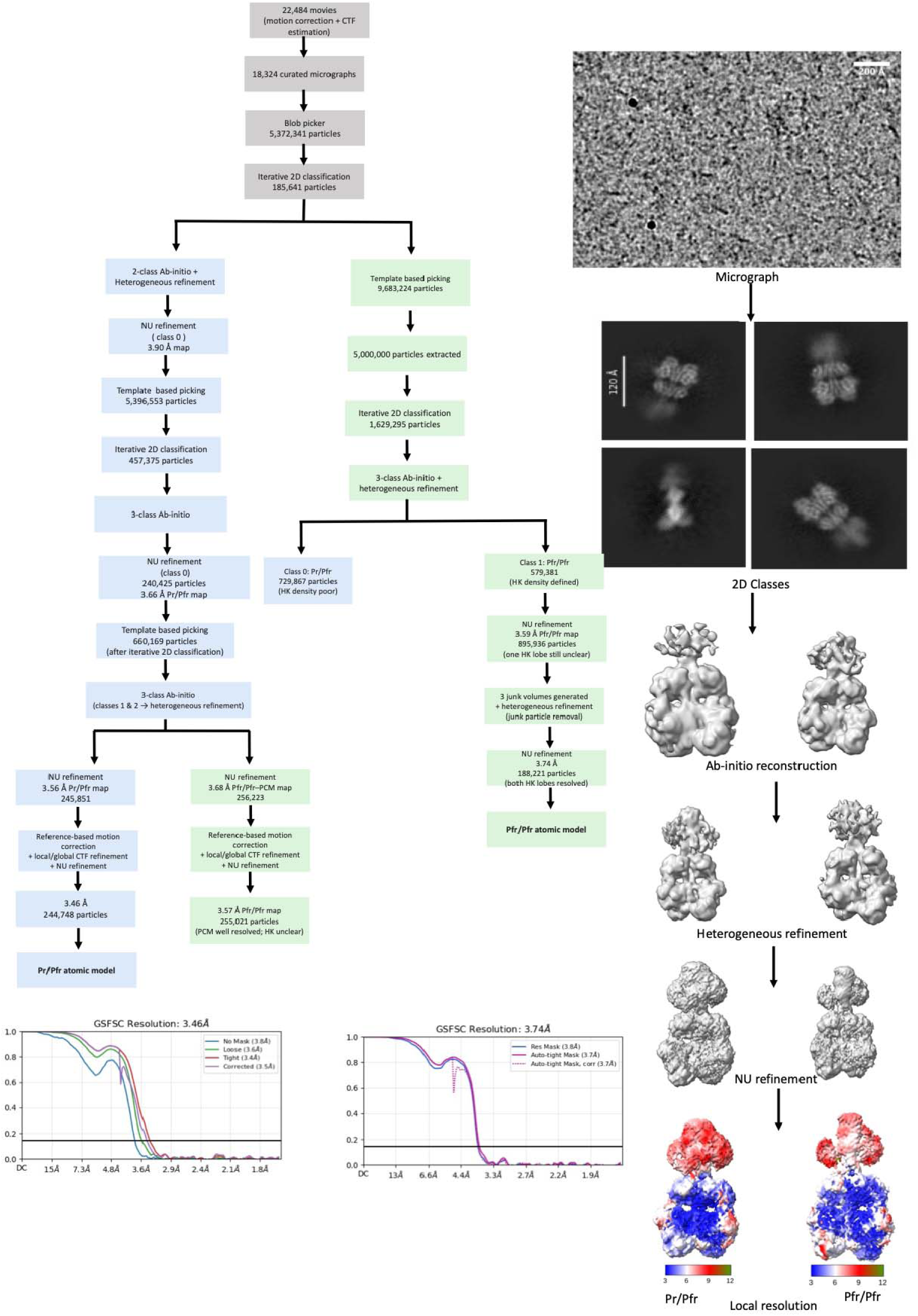
Flow chart for the processing of Cryo-EM data collected on the SaBphP Pr/Pfr heterodimer and the Pfr/Pfr homodimer species in the presence of ATP and the RR.

## Supplementary Tables

**Table S1.**
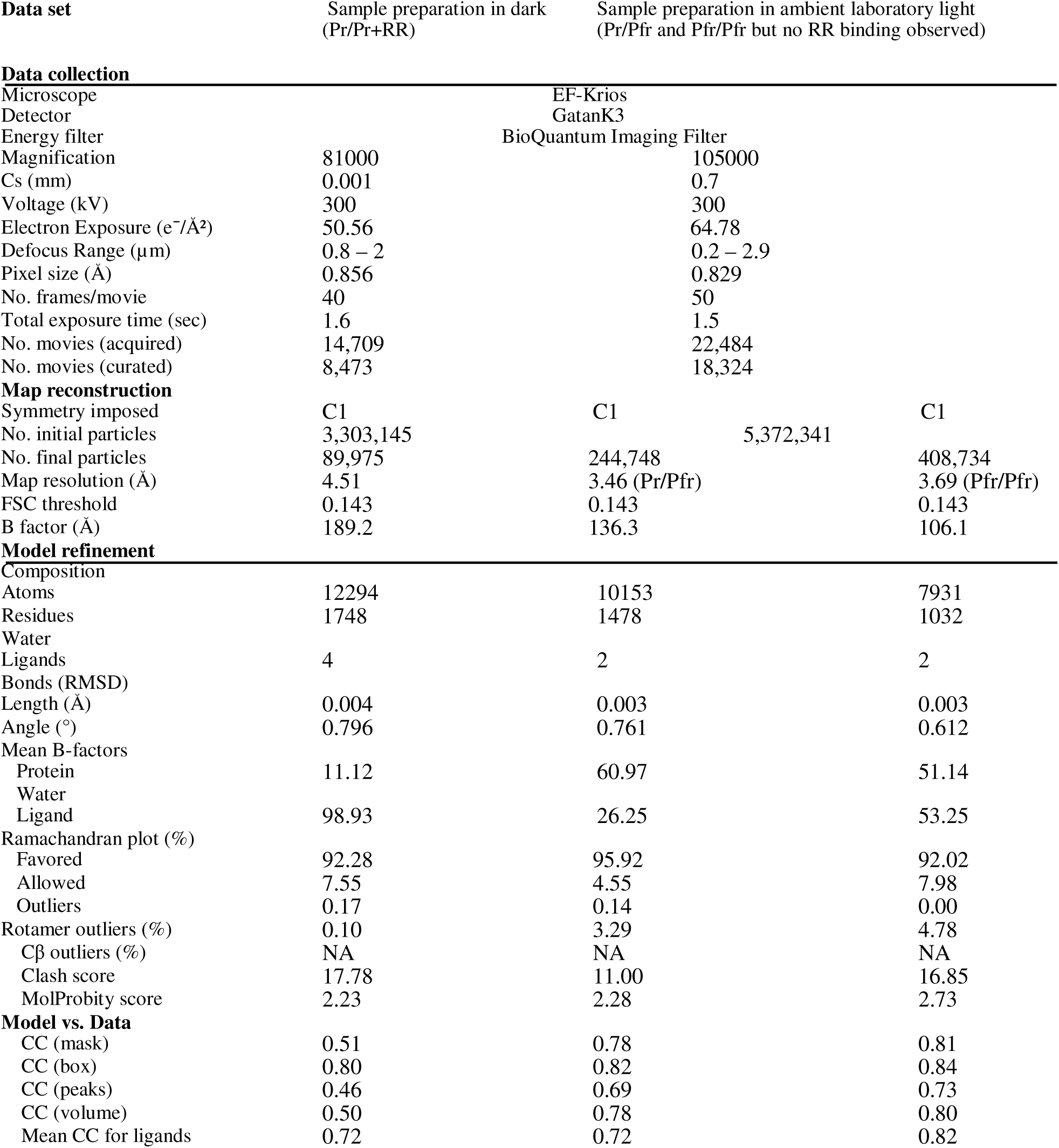
Statistics of cryo-EM data collection and low-resolution real space refinement.

| Data set | Sample preparation in dark<br>(Pr/Pr+RR) | Sample preparation in ambient laboratory light<br>(Pr/Pfr and Pfr/Pfr but no RR binding observed) |  |
| --- | --- | --- | --- |
| **Data collection** |  |  |  |
| Microscope |  | EF-Krios |  |
| Detector |  | GatanK3 |  |
| Energy filter |  | BioQuantum Imaging Filter |  |
| Magnification | 81000 |  | 105000 |
| Cs (mm) | 0.001 |  | 0.7 |
| Voltage (kV) | 300 |  | 300 |
| Electron Exposure (e<sup>-</sup>/Å<sup>2</sup>) | 50.56 |  | 64.78 |
| Defocus Range (µm) | 0.8 – 2 |  | 0.2 – 2.9 |
| Pixel size (Å) | 0.856 |  | 0.829 |
| No. frames/movie | 40 |  | 50 |
| Total exposure time (sec) | 1.6 |  | 1.5 |
| No. movies (acquired) | 14,709 |  | 22,484 |
| No. movies (curated) | 8,473 |  | 18,324 |
| **Map reconstruction** |  |  |  |
| Symmetry imposed | C1 | C1 | C1 |
| No. initial particles | 3,303,145 | 5,372,341 |  |
| No. final particles | 89,975 | 244,748 | 408,734 |
| Map resolution (Å) | 4.51 | 3.46 (Pr/Pfr) | 3.69 (Pfr/Pfr) |
| FSC threshold | 0.143 | 0.143 | 0.143 |
| B factor (Å) | 189.2 | 136.3 | 106.1 |
| **Model refinement** |  |  |  |
| Composition |  |  |  |
| Atoms | 12294 | 10153 | 7931 |
| Residues | 1748 | 1478 | 1032 |
| Water |  |  |  |
| Ligands | 4 | 2 | 2 |
| Bonds (RMSD) |  |  |  |
| Length (Å) | 0.004 | 0.003 | 0.003 |
| Angle (°) | 0.796 | 0.761 | 0.612 |
| Mean B-factors |  |  |  |
| Protein | 11.12 | 60.97 | 51.14 |
| Water |  |  |  |
| Ligand | 98.93 | 26.25 | 53.25 |
| Ramachandran plot (%) |  |  |  |
| Favored | 92.28 | 95.92 | 92.02 |
| Allowed | 7.55 | 4.55 | 7.98 |
| Outliers | 0.17 | 0.14 | 0.00 |
| Rotamer outliers (%) | 0.10 | 3.29 | 4.78 |
| Cβ outliers (%) | NA | NA | NA |
| Clash score | 17.78 | 11.00 | 16.85 |
| MolProbity score | 2.23 | 2.28 | 2.73 |
| **Model vs. Data** |  |  |  |
| CC (mask) | 0.51 | 0.78 | 0.81 |
| CC (box) | 0.80 | 0.82 | 0.84 |
| CC (peaks) | 0.46 | 0.69 | 0.73 |
| CC (volume) | 0.50 | 0.78 | 0.80 |
| Mean CC for ligands | 0.72 | 0.72 | 0.82 |

**Table S2.** X-ray Data collection and refinement statistics.

|  | RR_P43* |
| --- | --- |
| **Data collection** |  |
| Space group | P 4<sub>3</sub>2<sub>1</sub>2 |
| Cell dimensions |  |
| *a*, *b*, *c* (Å) | 86.08 86.08 38.50 |
| $\alpha$, $\beta$, $\gamma$ (°) | 90.0 90.0 90.0 |
| Resolution (Å) | 2.4 (2.45 – 2.4) |
| *R*<sub>merge</sub> [%] | 6.8 (0.82) |
| *cc*1/2 | 0.99 (0.84) |
| *I* / σ*I* | 12.0 (2.0) |
| Completeness (%) | 99.8 (100) |
| Redundancy | 6.3 (6.9) |
| **Refinement** |  |
| Resolution (Å) | 43.1 – 2.4 |
| No. reflections | 6884 |
| *R*<sub>work</sub> / *R*<sub>free</sub> [%] | 21.9/27.6 |
| No. atoms |  |
| Protein | 909 |
| Mg<sup>2+</sup> | 1 |
| Water | 6 |
| *B*-factors |  |
| Protein | 38.4 |
| Water | 46.1 |
| R.m.s. deviations |  |
| Bond lengths (Å) | 0.015 |
| Bond angles (°) | 2.0 |

\*one single crystal, bracket: highest resolution shell

## Supplementary Movies

The supplementary movies will be available once the manuscript is published in a peer reviewed journal.

**Movie S1: The SaBphP-RR complex.** The SaBphP_Pr/Pr_-RR complex is rotated by 360° about the y-axis to visualize the interaction of the HK with the RR in more detail. The cryo-EM density that covers the HK region including the 4-helix bundle and part of the linker helices as well as the RR is shown in transparent gray. The SaBphP2 protomer A is shown in blue that of B in red. The chromophores are depicted in their respective protomers in opposite colors. RR subunit A is shown in yellow and subunit B in green.

**Movie S2: The 4-helix bundle at improved resolution.** Caused by the binding of the RR, the local resolution in the region of the SaBphP 4-helix bundle increases relative to the free BphP. The cryo-EM density (light blue) unambiguously shows to which BphP protomer the HK lobes are bound (green: SaBphP protomer A, red: SaBphP2, protomer B).

**Movie S3: Binding and dissociation of the RR to SaBphP2**. The movie was generated by a morph across 4 different structures: The structures are: (1) the SaBphP2-Pr/Pr homodimer without the RR, (2) the Pr/Pr homodimer with the RR bound (the SaBphP2-RR complex), (3) the SaBphP2-Pr/Pfr heterodimer and (4) the Pfr/Pfr homodimer. Structures (2) to (4) were determined in the presence of ATP and the RR. Since the movie is a morph that extrapolates between observed structures, structural details during the transitions between the structures are mostly incorrect. However, the movie provides a visual representation of the overall progress of the reaction, which maps the TCS pathway of the light activated receptor kinase. The red flash indicates irradiation by red light to trigger the BV *Z* to *E* isomerization in the Pr/Pr homodimer that results in the 180° HK rotation and the RR release.

## References

1. Y. Xie, J. Li, Y. Ding, X. Shao, Y. Sun, F. Xie, S. Liu, S. Tang, X. Deng, An atlas of bacterial two-component systems reveals function and plasticity in signal transduction. Cell reports 41, 111502 (2022), 10.1016/j.celrep.2022.111502,

2. A. M. Stock, V. L. Robinson, P. N. Goudreau, Two-component signal transduction. Annu Rev Biochem 69, 183–215 (2000), 10.1146/annurev.biochem.69.1.183,

3. R. Gao, S. Bouillet, A. M. Stock, Structural Basis of Response Regulator Function. Annu Rev Microbiol 73, 175–197 (2019), 10.1146/annurev-micro-020518-115931,

4. J. T. Lazar, J. J. Tabor, Bacterial two-component systems as sensors for synthetic biology applications. Curr Opin Syst Biol 28, (2021), 10.1016/j.coisb.2021.100398, PMC8670732.

5. A. Buschiazzo, F. Trajtenberg, Two-Component Sensing and Regulation: How Do Histidine Kinases Talk with Response Regulators at the Molecular Level? Annu Rev Microbiol 73, 507–528 (2019), 10.1146/annurev-micro-091018-054627,

6. H. Hirakawa, J. Kurushima, Y. Hashimoto, H. Tomita, Progress Overview of Bacterial Two-Component Regulatory Systems as Potential Targets for Antimicrobial Chemotherapy. Antibiotics (Basel) 9, (2020), 10.3390/antibiotics9100635, PMC7598275.

7. A. M. Stock, J. M. Mottonen, J. B. Stock, C. E. Schutt, Three-dimensional structure of CheY, the response regulator of bacterial chemotaxis. Nature 337, 745–749 (1989), 10.1038/337745a0,

8. P. Casino, V. Rubio, A. Marina, Structural Insight into Partner Specificity and Phosphoryl Transfer in Two-Component Signal Transduction. Cell 139, 325–336 (2009), 10.1016/j.cell.2009.08.032,

9. S. Yamada, H. Sugimoto, M. Kobayashi, A. Ohno, H. Nakamura, Y. Shiro, Structure of PAS-linked histidine kinase and the response regulator complex. Structure 17, 1333–1344 (2009), 10.1016/j.str.2009.07.016,

10. F. Trajtenberg, J. A. Imelio, M. R. Machado, N. Larrieux, M. A. Marti, G. Obal, A. E. Mechaly, A. Buschiazzo, Regulation of signaling directionality revealed by 3D snapshots of a kinase:regulator complex in action. eLife 5, (2016), 10.7554/eLife.21422, PMC5231405.

11. J. C. Sanchez, M. Carrillo, S. Pandey, M. Noda, L. Aldama, D. Feliz, E. Claesson, W. Y. Wahlgren, G. Tracy, P. Duong, A. C. Nugent, A. Field, V. Srajer, C. Kupitz, S. Iwata, E. Nango, R. Tanaka, T. Tanaka, L. Fangjia, K. Tono, S. Owada, S. Westenhoff, M. Schmidt, E. A. Stojkovic, High-resolution crystal structures of amyxobacterial phytochrome at cryo and roomtemperatures. Struct Dyn-Us 6, (2019), 10.1063/1.5120527,

12. P. Karki, D. Menendez, W. Budell, S. Dangi, C. Hernandez, J. Mendez, S. Muniyappan, S. Basu, P. Schwander, T. N. Malla, E. A. Stojkovic, M. Schmidt, Structures of myxobacterial phytochrome revealed by cryo-EM using the Spotiton technique and with x-ray crystallography. Struct Dyn 12, 034701 (2025), 10.1063/4.0000301, PMC12048173.

13. P. Karki, W. C. Budell, R. Kannachel, D. Menendez, C. Hernandez, J. H. Mendez, S. Kanou, M. Singh, C. Slavov, P. Schwander, T. N. Malla, E. A. Stojkovic, M. Schmidt, A bacteriophytochrome Pr/Pfr heterodimer studied through single-particle time-resolved cryo-electron microscopy. Commun Chem, (2026), 10.1038/s42004-026-02084-6,

14. S. Huntley, N. Hamann, S. Wegener-Feldbrugge, A. Treuner-Lange, M. Kube, R. Reinhardt, S. Klages, R. Muller, C. M. Ronning, W. C. Nierman, L. Sogaard-Andersen, Comparative genomic analysis of fruiting body formation in Myxococcales. Mol Biol Evol 28, 1083–1097 (2011), 10.1093/molbev/msq292,

15. N. C. Woitowich, A. S. Halavaty, P. Waltz, C. Kupitz, J. Valera, G. Tracy, K. D. Gallagher, E. Claesson, T. Nakane, S. Pandey, G. Nelson, R. Tanaka, E. Nango, E. Mizohata, S. Owada, K. Tono, Y. Joti, A. C. Nugent, H. Patel, A. Mapara, J. Hopkins, P. Duong, D. Bizhga, S. E. Kovaleva, R. St Peter, C. N. Hernandez, W. B. Ozarowski, S. Roy-Chowdhuri, J. H. Yang, P. Edlund, H. Takala, J. Ihalainen, J. Brayshaw, T. Norwood, I. Poudyal, P. Fromme, J. C. H. Spence, K. Moffat, S. Westenhoff, M. Schmidt, E. A. Stojkovic, Structural basis for light control of cell development revealed by crystal structures of a myxobacterial phytochrome. IUCrJ 5, 619–634 (2018), 10.1107/S2052252518010631, PMC6126659.

16. P. Lapuhs, E. Heinrich, R. Garcia, A. Goes, N. Frank, L. Bollenbach, V. Stibane, T. Kuhn, M. Koch, A. K. Kiemer, R. Muller, K. Fuhrmann, G. Fuhrmann, The inherent antibiotic activity of myxobacteria-derived autofluorescent outer membrane vesicles is switched on and off by light stimulation. Nanoscale 14, 17534–17542 (2022), 10.1039/d2nr02743g,

17. J. Han, M. Shi, X. Dou, W. Pan, D. Ma, M. Luo, B. Fu, Cystobacter fuscus HM-E: a novel biocontrol agent against cotton Verticillium wilt. Front Microbiol 16, 1555523 (2025), 10.3389/fmicb.2025.1555523, PMC11936981.

18. L. Zhang, X. Ye, Z. Zhang, Y. Li, Z. Cui, D. Wall, Z. Li, Bacterial predator-prey interactions and their potentials in combating plant pathogens. Microbiol Res 310, 128559 (2026), 10.1016/j.micres.2026.128559,

19. M. Vasse, G. J. Velicer, Predation in microbial communities: gradients of nutritive killing. Nat Rev Microbiol, (2026), 10.1038/s41579-026-01299-7,

20. M. E. Auldridge, K. T. Forest, Bacterial phytochromes: More than meets the light. Crit Rev Biochem Mol 46, 67–88 (2011), Doi 10.3109/10409238.2010.546389,

21. T. N. Malla, C. Hernandez, S. Muniyappan, D. Menendez, D. Bizhga, J. H. Mendez, P. Schwander, E. A. Stojkovic, M. Schmidt, Photoreception and signaling in bacterial phytochrome revealed by single-particle cryo-EM. Sci Adv 10, eadq0653 (2024), 10.1126/sciadv.adq0653, PMC11313861.

22. A. W. Baker, K. A. Satyshur, K. T. Forest. (2016), doi.org/10.2210/pdb5BRJ/pdb, PMCID.

23. 23. B. Karniol, R. D. Vierstra, The pair of bacteriophytochromes from Agrobacterium tumefaciens are histidine kinases with opposing photobiological properties. Proceedings of the National Academy of Sciences of the United States of America 100, 2807–2812 (2003), 10.1073/pnas.0437914100, PMC151422.

24. X. Yang, K. Moffat. (2015), doi.org/10.2210/pdb4ZYL/pdb, PMCID.

25. 25. E. Multamaki, R. Nanekar, D. Morozov, T. Lievonen, D. Golonka, W. Y. Wahlgren, B. Stucki-Buchli, J. Rossi, V. P. Hytonen, S. Westenhoff, J. A. Ihalainen, A. Moglich, H. Takala, Comparative analysis of two paradigm bacteriophytochromes reveals opposite functionalities in two-component signaling. Nat Commun 12, 4394 (2021), 10.1038/s41467-021-24676-7, PMC8292422.

26. I. Baikalov, I. Schroder, M. Kaczor-Grzeskowiak, K. Grzeskowiak, R. P. Gunsalus, R. E. Dickerson, Structure of the Escherichia coli response regulator NarL. Biochemistry 35, 11053–11061 (1996), 10.1021/bi960919o,

27. K. Volz, P. Matsumura, Crystal structure of Escherichia coli CheY refined at 1.7-A resolution. The Journal of biological chemistry 266, 15511–15519 (1991), 10.2210/pdb3chy/pdb,

28. D. R. Buckler, Y. Zhou, A. M. Stock, Evidence of intradomain and interdomain flexibility in an OmpR/PhoB homolog from Thermotoga maritima. Structure 10, 153–164 (2002), 10.1016/s0969-2126(01)00706-7,

29. A. W. Baker, K. A. Satyshur, N. Moreno Morales, K. T. Forest, Arm-in-Arm Response Regulator Dimers Promote Intermolecular Signal Transduction. J Bacteriol 198, 1218–1229 (2016), 10.1128/JB.00872-15, PMC4859592.

30. A. W. Baker, K. A. Satyshur, K. T. Forest. (2016), doi.org/10.2210/pdb5IC5/pdb, PMCID.

31. 31. Y. Liu, Z. Q. Gao, Z. She, K. Qu, W. J. Wang, E. V. Shtykova, J. H. Xu, C. N. Ji, Y. H. Dong, The structural basis of the response regulator DrRRA from Deinococcus radiodurans. Biochemical and biophysical research communications 417, 1206–1212 (2012), 10.1016/j.bbrc.2011.12.110,

32. S. Y. Lee, H. S. Cho, J. G. Pelton, D. Yan, R. K. Henderson, D. S. King, L. Huang, S. Kustu, E. A. Berry, D. E. Wemmer, Crystal structure of an activated response regulator bound to its target. Nat Struct Biol 8, 52–56 (2001), 10.1038/83053,

33. P. R. Thompson, P. A. Cole, Probing the mechanism of enzymatic phosphoryl transfer with a chemical trick. Proceedings of the National Academy of Sciences of the United States of America 98, 8170–8171 (2001), 10.1073/pnas.161273998, PMC37416.

34. S. Ganguli, H. Wang, P. Matsumara, K. Volz, Uncoupled phosphorylation and activation of bacterial chemotaxis. The Journal of biological chemistry 270, 17386–17393 (1995),

35. S. Y. Lee, H. S. Cho, J. G. Pelton, D. Yan, E. A. Berry, D. E. Wemmer, Crystal structure of activated CheY. Comparison with other activated receiver domains. The Journal of biological chemistry 276, 16425–16431 (2001), 10.1074/jbc.M101002200,

36. I. Kumarapperuma, I. P. Tom, S. Bandara, S. Montano, X. Yang, Mode of autophosphorylation in bacteriophytochromes RpBphP2 and RpBphP3. Photochemical & photobiological sciences : Official journal of the European Photochemistry Association and the European Society for Photobiology 22, 1257–1266 (2023), 10.1007/s43630-023-00366-9, PMC10619329.

37. Y. Yang, M. Inouye, Intermolecular complementation between two defective mutant signal-transducing receptors of Escherichia coli. Proceedings of the National Academy of Sciences of the United States of America 88, 11057–11061 (1991), 10.1073/pnas.88.24.11057, PMC53072.

38. E. G. Ninfa, M. R. Atkinson, E. S. Kamberov, A. J. Ninfa, Mechanism of autophosphorylation of Escherichia coli nitrogen regulator II (NRII or NtrB): trans-phosphorylation between subunits. J Bacteriol 175, 7024–7032 (1993), 10.1128/jb.175.21.7024-7032.1993, PMC206830.

39. K. B. Graniczkowska, D. Bizhga, M. Noda, V. Leon, N. Saraf, D. Feliz, G. Sharma, A. C. Nugent, M. Singer, E. A. Stojkovic, Photomorphogenesis of Myxococcus macrosporus: New Insights for Light-Regulation of Cell Development. Photoch Photobio Sci, 14 (2024), 10.1007/s43630-024-00635-1,

40. T. N. Malla, L. Aldama, V. Leon, D. Feliz, H. Hu, I. Thomas, A. Cellini, W. Y. Wahlgren, A. Nimmrich, S. Botha, R. Sierra, M. S. Hunter, F. Poitevin, S. Lisova, A. Batyuk, G. Gate, R. Jernigan, C. J. Kupitz, P. Maj, P. Meszaros, M. Kurttila, L. Monrroy, F. Luo, S. Owada, J. Kang, C. Slavov, M. Maj, C. Gautier, M. Kashipathy, A. Tolstikova, V. Mariani, A. Barty, F. Moss, P. Schwander, H. Liu, S. Boutet, P. Fromme, H. Takala, J. A. Ihalainen, U. Weierstall, S. Westenhoff, E. A. Stojkovic, M. Schmidt, Observation of early events in the photoactivation of Myxobacterial phytochrome using time-resolved serial femtosecond crystallography. Commun Chem 8, 183 (2025), 10.1038/s42004-025-01578-z, PMC12162953.

41. M. Carrillo, S. Pandey, J. Sanchez, M. Noda, I. Poudyal, L. Aldama, T. N. Malla, E. Claesson, W. Y. Wahlgren, D. Feliz, V. Srajer, M. Maj, L. Castillon, S. Iwata, E. Nango, R. Tanaka, T. Tanaka, L. Fangjia, K. Tono, S. Owada, S. Westenhoff, E. A. Stojkovic, M. Schmidt, High-resolution crystal structures of transient intermediates in the phytochrome photocycle. Structure, (2021), 10.1016/j.str.2021.03.004,

42. J. M. Skerker, B. S. Perchuk, A. Siryaporn, E. A. Lubin, O. Ashenberg, M. Goulian, M. T. Laub, Rewiring the specificity of two-component signal transduction systems. Cell 133, 1043–1054 (2008), 10.1016/j.cell.2008.04.040, PMC2453690.

43. C. Liu, D. Sun, J. Zhu, W. Liu, Two-Component Signal Transduction Systems: A Major Strategy for Connecting Input Stimuli to Biofilm Formation. Front Microbiol 9, 3279 (2018), 10.3389/fmicb.2018.03279, PMC6335343.

44. R. D. Bardgett, W. H. van der Putten, Belowground biodiversity and ecosystem functioning. Nature 515, 505–511 (2014), 10.1038/nature13855,

45. H. M. Berman, T. Battistuz, T. N. Bhat, W. F. Bluhm, P. E. Bourne, K. Burkhardt, Z. Feng, G. L. Gilliland, L. Iype, S. Jain, P. Fagan, J. Marvin, D. Padilla, V. Ravichandran, B. Schneider, N. Thanki, H. Weissig, J. D. Westbrook, C. Zardecki, The Protein Data Bank. Acta crystallographica. Section D, Biological crystallography 58, 899–907 (2002),

46. wwPDB-consortium, EMDB-the Electron Microscopy Data Bank. Nucleic Acids Res 52, D456–D465 (2024), 10.1093/nar/gkad1019, PMC10767987.

47. M. Kostovski, A. Oren, Proposal of Allogeochorda gen. nov., Allogeochorda subterranea comb. nov., Allostigmatella gen. nov. and four new combinations and Allosphingorhabdus gen. nov. and Allosphingorhabdus lutea comb. nov. as replacement names for the illegitimate genus names Geochorda, Stigmatella and Alterisphingorhabdus and their species. Int J Syst Evol Microbiol 76, (2026), 10.1099/ijsem.0.007208, PMC13281875.

48. A. Punjani, J. L. Rubinstein, D. J. Fleet, M. A. Brubaker, cryoSPARC: algorithms for rapid unsupervised cryo-EM structure determination. Nature methods 14, 290–296 (2017), 10.1038/nmeth.4169,

49. E. F. Pettersen, T. D. Goddard, C. C. Huang, E. C. Meng, G. S. Couch, T. I. Croll, J. H. Morris, T. E. Ferrin, UCSF ChimeraX: Structure visualization for researchers, educators, and developers. Protein Sci 30, 70–82 (2021), 10.1002/pro.3943, PMC7737788.

50. D. Liebschner, P. V. Afonine, M. L. Baker, G. Bunkoczi, V. B. Chen, T. I. Croll, B. Hintze, L. W. Hung, S. Jain, A. J. McCoy, N. W. Moriarty, R. D. Oeffner, B. K. Poon, M. G. Prisant, R. J. Read, J. S. Richardson, D. C. Richardson, M. D. Sammito, O. V. Sobolev, D. H. Stockwell, T. C. Terwilliger, A. G. Urzhumtsev, L. L. Videau, C. J. Williams, P. D. Adams, Macromolecular structure determination using X-rays, neutrons and electrons: recent developments in Phenix. Acta Crystallogr D Struct Biol 75, 861–877 (2019), 10.1107/S2059798319011471, PMC6778852.

51. T. G. Battye, L. Kontogiannis, O. Johnson, H. R. Powell, A. G. Leslie, iMOSFLM: a new graphical interface for diffraction-image processing with MOSFLM. Acta crystallographica. Section D, Biological crystallography 67, 271–281 (2011), 10.1107/S0907444910048675, PMC3069742.

52. J. Abramson, J. Adler, J. Dunger, R. Evans, T. Green, A. Pritzel, O. Ronneberger, L. Willmore, A. J. Ballard, J. Bambrick, S. W. Bodenstein, D. A. Evans, C. C. Hung, M. O’Neill, D. Reiman, K. Tunyasuvunakool, Z. Wu, A. Zemgulyte, E. Arvaniti, C. Beattie, O. Bertolli, A. Bridgland, A. Cherepanov, M. Congreve, A. I. Cowen-Rivers, A. Cowie, M. Figurnov, F. B. Fuchs, H. Gladman, R. Jain, Y. A. Khan, C. M. R. Low, K. Perlin, A. Potapenko, P. Savy, S. Singh, A. Stecula, A. Thillaisundaram, C. Tong, S. Yakneen, E. D. Zhong, M. Zielinski, A. Zidek, V. Bapst, P. Kohli, M. Jaderberg, D. Hassabis, J. M. Jumper, Accurate structure prediction of biomolecular interactions with AlphaFold 3. Nature 630, 493–500 (2024), 10.1038/s41586-024-07487-w, PMC11168924.

